# Tumor context determines *ARID1A* effects on gastric cancer immunity

**DOI:** 10.64898/2026.08.18.745460

**Authors:** Shu Xiao, Ryan T. Heslin, Morgan F. Pettigrew, John D. Karalis, Nafeesah Fatimah, Shao-Po Huang, Verena Cao, Esther Burns, Lynn Y. Kwon, Ibrahim Nassour, Skylar L. Nahi, Hsien Tsung Lai, Changjin Hong, Tae Hyun Hwang, Isaac S. Chan, Suntrea T.G. Hammer, Hao Zhu, Sam C. Wang

**Author notes:** These authors contributed equally (Shu Xiao, Ryan T. Heslin, Morgan F. Pettigrew, John D. Karalis). **Corresponding Author:** Sam C. Wang, MD, Department of Surgery, University of Texas Southwestern Medical Center, 5323 Harry Hines Boulevard, Dallas, Texas 75390. The authors declare no relevant conflicts of interest.

## Abstract

The role of *ARID1A* in cancer immune evasion remains uncertain, with prior studies reaching opposing conclusions. In addition, previous work has shown that the role of *ARID1A* in cell-autonomous tumorigenesis is context-dependent. Using isogenic murine gastric cancer models, we found that *in vivo Arid1a* loss in an autochthonous genetically engineered mouse model of gastric cancer conferred T cell-dependent immune evasion, while *in vitro* deletion did not. Mechanistically, tumor *Arid1a* loss reprogrammed the tumor microenvironment into an immune desert through suppression of GM-CSF secretion and interferon-γ responsiveness. These changes were not observed when *Arid1a* was deleted *in vitro*. In human gastric cancer, an immune-cold phenotype was restricted to *ARID1A* mutants in the genomically stable subtype, while *ARID1A* loss in the chromosomal instability subtype was associated with variable immune profiles. These results demonstrate that tumor *ARID1A* loss does not intrinsically confer pro- or anti-tumor immune properties and instead is determined by tissue context.

## BACKGROUND

*ARID1A* is a component of the SWI/SNF chromatin remodeling complex and is one of the most commonly mutated genes in cancer.^1^ *ARID1A* is generally considered a tumor suppressor; however, we have previously shown that *ARID1A* is permissive for cancer formation in normal liver tissue, demonstrating that *ARID1A* has context-specific effects.^2^

Whether *ARID1A* induces pro- or anti-tumor immunity is uncertain due to conflicting reports. Some groups have shown that cancers with loss-of-function *ARID1A* mutations become more immunogenic and responsive to immunotherapies.^3–5^ However, others report that *ARID1A* loss leads to pro-tumorigenic immune effects.^6–8^ Similarly, there have been conflicting reports of how tumor *ARID1A* mutations correlate to immunotherapy response in the clinic.^9–11^ Delineating the immune effects of *ARID1A* mutations is clinically necessary as *ARID1A* mutation status is being tested as a stratification biomarker for immunotherapy use (ClinicalTrials.gov: NCT04957615, NCT06518564, NCT04953104, NCT04065269). Additionally, the introduction of SWI/SNF-targeted inhibitors in early-phase clinical trials further emphasizes the need for a deeper understanding of the mechanisms that link tumor *ARID1A* status to tumor immunity.^12^

Tumor cell immunogenicity is shaped, in part, by host contexts that exert selection pressures that “sculpt” cancer cells. In this process, known as immunoediting, tumor clones that elude immune clearance persist until they acquire the capability to completely escape immune control, at which point the cancer progresses.^13,14^ In this study, we compared gastric cancers in which tumor *ARID1A* loss occurred under different contexts. We established gastric cancer organoids in which *Arid1a* was deleted *in vitro* and compared them to an autochthonous genetically engineered mouse model in which *Arid1a* was deleted *in vivo*. We found that *Arid1a* loss promoted immune evasion only in the *in vivo* deletion model, through altered GM-CSF and IFNGR1 expression that blunted the initiation of the cancer immunity cycle. These alterations were absent when *Arid1a* was deleted *in vitro*. We next confirmed the context-dependency of *ARID1A* function in human patients. We analyzed The Cancer Genome Atlas gastric adenocarcinoma cohort and found that the association of *ARID1A* mutation to immunogenicity differed based on molecular subtype. Together, our findings indicate that the immune effect of tumor *ARID1A* loss is context-dependent and suggest that divergent results from previous studies on cancer immune effects of *ARID1A* may be due to the different molecular and host contexts that each study utilized.

## RESULTS

### *In vitro Arid1a* deletion in gastric cancer tumoroids promoted tumor growth but not immune evasion

*TP53* and *CDH1* are commonly altered tumor suppressor genes in gastric cancer.^15^ We generated gastric cancer organoids (tumoroids) from mice with the genetic backgrounds of *Trp53^flox/flox^; Cdh1^flox/flox^; Rosa26^LSL-eYFP/LSL-eYFP^* and either *Arid1a*^+/+^ or *Arid1a*^flox/flox^ (**Figure 1A**).^16^ Once the tumoroids were established, adenovirus carrying *Cre* recombinase was added to produce *in vitro* recombination (*in-vitro*-R) of the floxed alleles to result in *Trp53* and *Cdh1* deletion, *eYFP* expression, and *Arid1a* deletion in *Arid1a^flox/flox^* tumoroids (**Figure 1B)**. We refer to the tumoroid lines as *in-vitro*-R *Arid1a* wild-type (WT) and *in-vitro*-R *Arid1a* knockout (KO). The tumoroids were placed under nutlin-3a selection to enrich for *Trp53*-null cells.^17^ We confirmed the expected deletion of *Trp53* and *Cdh1* (**Figure 1C; Supplemental Figure 1**) and confirmed all tumoroids expressed eYFP (**Figure 1D**).

**Figure 1.**
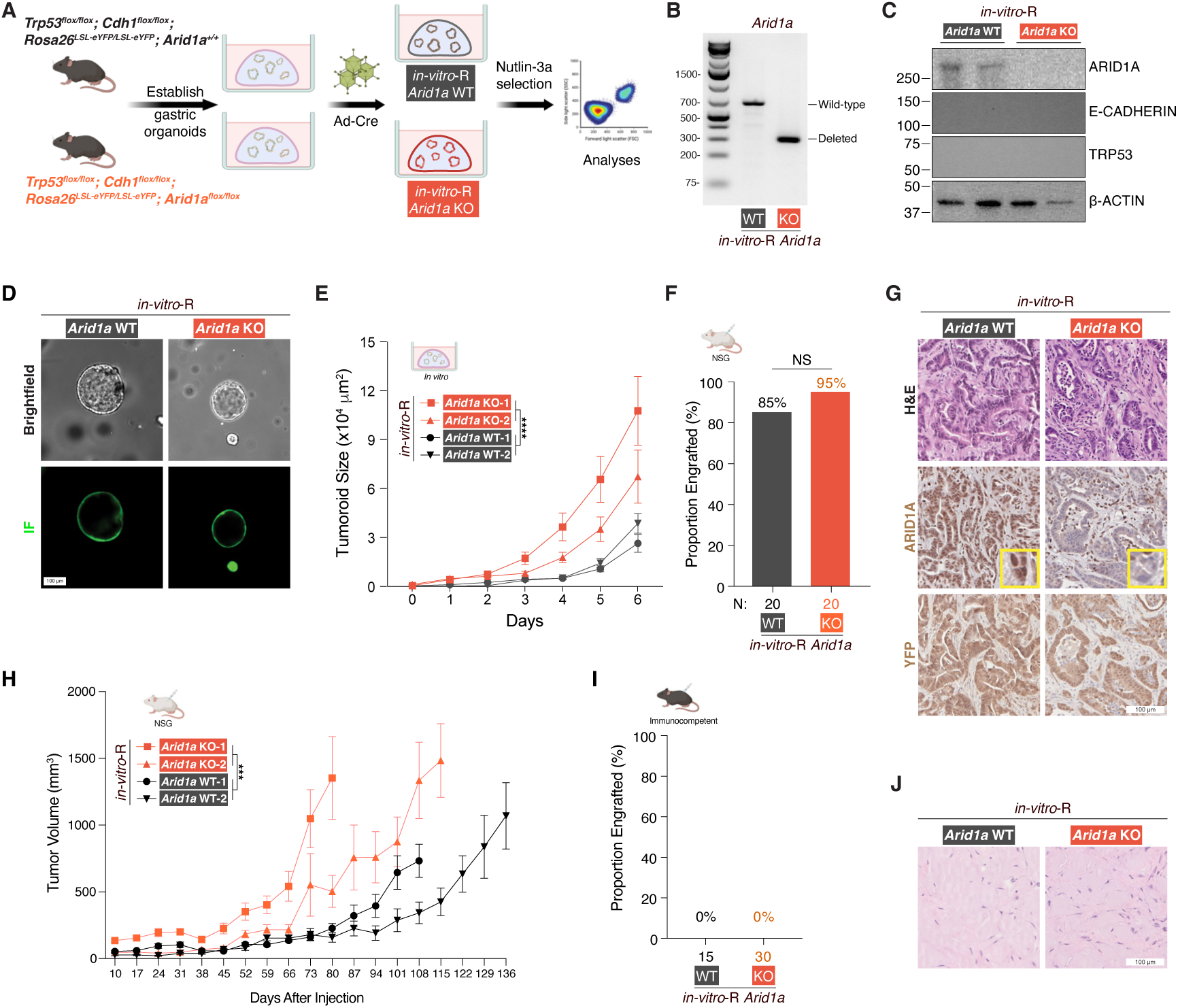
*In vitro Arid1a* deletion in gastric cancer tumoroids promoted tumor growth but not immune evasion. **(A)** Schematic of tumoroid generation with *in vitro Arid1a* deletion. These tumoroids are termed *in-vitro*-R *Arid1a* wild-type (WT) or *in-vitro*-R *Arid1a* knockout (KO). **(B)** Representative genotyping PCR of *in-vitro*-R *Arid1a* WT and KO tumoroids after Cre recombination. **(C)** Western blots for ARID1A, E-CADHERIN, TRP53, and β-ACTIN from *in-vitro*-R *Arid1a* WT and KO tumoroids after Cre recombination. See also Supplemental Figure 1. **(D)** Representative brightfield (top) and immunofluorescence (IF; bottom) images of *in-vitro*-R *Arid1a* WT and KO tumoroids after Cre recombination. **(E)** Tumoroid size over time measured by live imaging for *in-vitro-*R *Arid1a* WT and KO lines. Data shown as mean ± SEM. Statistical comparison of pooled WT (N = 184) vs pooled KO (N = 140) tumoroids through day 6 by mixed-effects model (REML) with Geisser-Greenhouse correction; asterisks denote the time × genotype interaction. **(F)** Proportion of NSG mice that developed engrafted tumors following flank injection of *in-vitro*-R *Arid1a* WT or KO tumoroids. Proportions compared via Fisher’s exact test. **(G)** Representative H&E (top), ARID1A (middle), and YFP (bottom) expression in tumors generated from NSG mice injected with *in-vitro*-R *Arid1a* WT or KO tumoroids. **(H)** Tumor volume over time following flank injection of *in-vitro*-R *Arid1a* WT and KO tumoroids into NSG mice. Data shown as mean ± SEM. Statistical comparison of pooled WT (N = 20) vs pooled KO (N = 22) genotypes through day 80 (last shared timepoint) by mixed-effects model (REML) with Geisser-Greenhouse correction; asterisks denote the time × genotype interaction. **(I)** Proportion of syngeneic immunocompetent mice that developed engrafted tumors following flank injection of *in-vitro*-R *Arid1a* WT or KO tumoroids. Proportions compared via Fisher’s exact test. **(J)** Representative H&E of tissue from syngeneic immunocompetent mice 4 months after flank injection of *in-vitro*-R *Arid1a* WT or KO tumoroids, showing collagenized scar without residual carcinoma. NS, non-significant, *\** P < 0.05, ** P < 0.01, *** P < 0.001, **** P < 0.0001.

We first compared the growth kinetics of the tumoroids and found *in-vitro*-R *Arid1a* KO lines grew faster than the *in-vitro*-R *Arid1a* WT lines, consistent with a tumor suppressive function for *Arid1a* (P < 0.0001; **Figure 1E**). To confirm this finding in an *in vivo* setting, we injected the tumoroids into the flanks of NOD.Cg-*Prkdc^scid^ Il2rg^tm1Wjl^*/SzJ (NSG) mice and found that both *in-vitro-*R *Arid1a* WT and KO tumoroids engrafted robustly, which we defined as tumors reaching at least 200 mm^3^ followed by continuous growth (**Figure 1F**). The engrafted tumors appeared histologically similar to gastric adenocarcinoma, had the expected ARID1A expression based on the genotype, and expressed YFP (**Figure 1G**). Consistent with the *in vitro* growth assay, the engrafted *in-vitro-*R *Arid1a* KO tumors grew faster than the *Arid1a* WT tumors (P < 0.001; **Figure 1H**).

To determine whether tumor *Arid1a* status affected their interaction with the host immune system, we injected *in-vitro*-R *Arid1a* WT and KO tumoroids into syngeneic littermate controls of the mice from which we generated the tumoroids. We found neither tumoroid genotype was able to engraft effectively and produce a growing tumor in these immunocompetent hosts (**Figure 1I**). Instead, for both genotypes, the tumor either completely regressed or remained very small and stable in size. On H&E, the masses that were present 4 months post-injection consisted of only mature collagenized scar without evidence of residual carcinoma. Some scarred areas were surrounded by mononuclear inflammatory cells (**Figure 1J**). These data suggested that while *Arid1a* loss promoted growth in an immunodeficient setting, *Arid1a* loss did not support immune evasion. Both genotypes were effectively cleared by the immune system, which may be due to the antigenicity that the eYFP produced.^18^

### *In vivo* tumor *Arid1a* loss in genetically engineered mice promoted aggressive features and shortened survival

We next tested the role of *Arid1a* on gastric cancer biology in an autochthonous transgenic mouse model of gastric cancer.^19^ We generated *Atp4b-Cre; Trp53^flox/flox^; Cdh1^flox/flox^; Rosa26^LSL-eYFP/LSL-eYFP^*mice that were *Arid1a^+/+^* (WT), *Arid1a^+/flox^*(Het), or *Arid1a^flox/flox^* (KO; **Figure 2A**). *Atp4b* is expressed only in gastric parietal lineage cells, which differentiate into the acid-producing cells in the stomach.^19,20^ These mice were isogenic to the mice from which *in-vitro-*R *Arid1a* WT and KO tumoroids were derived. We refer to this system that had *in vivo* recombination of floxed alleles as *in-vivo*-R.

**Figure 2.**
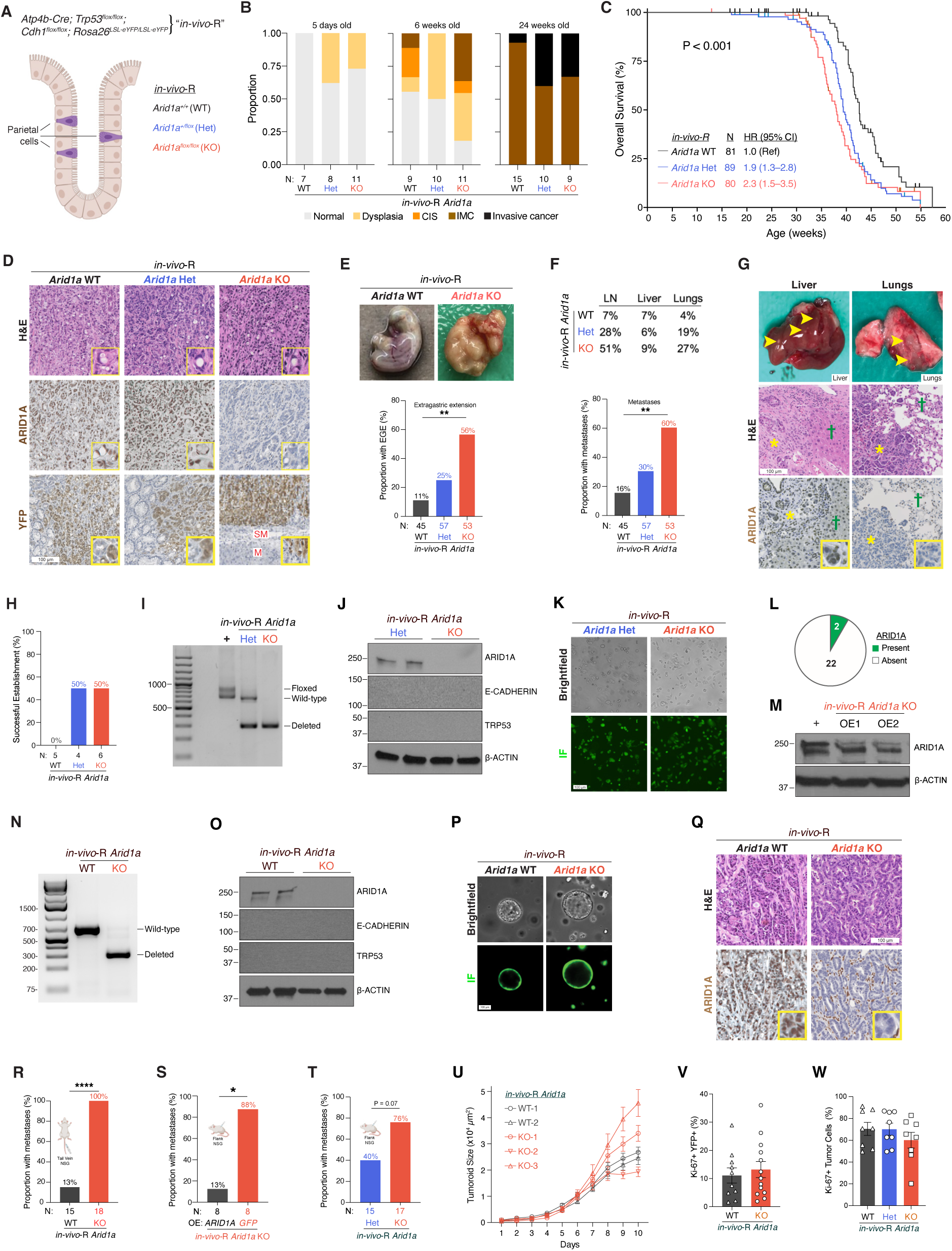
*In vivo* tumor *Arid1a* loss in a genetically engineered mouse model promoted aggressive disease features and shortened survival. **(A)** Schematic of transgenic mouse model of gastric cancer, which had *in vivo* recombination (*in-vivo*-R) of all the floxed alleles, leading to autochthonous deletion of *Trp53, Cdh1*, and *Arid1a*. *Atp4b-Cre*; *Trp53*^flox/flox^; *Cdh1*^flox/flox^; *Rosa26*^LSL-eYFP/LSL-eYFP^ mice were generated with *Arid1a*^+/+^ (*in-vivo*-R *Arid1a* WT), *Arid1a*^+/flox^ (*in-vivo*-R *Arid1a* Het), or *Arid1a*^flox/flox^ (*in-vivo*-R *Arid1a* KO) backgrounds. *Atp4b-Cre* drives recombination selectively in gastric parietal cells. **(B)** Histologic assessment of gastric mucosa from *in-vivo*-R *Arid1a* WT, Het, and KO mice at 5 days, 6 weeks, and 24 weeks of age. Stacked bars show the proportion of mice with normal mucosa, dysplasia, carcinoma in situ (CIS), intramucosal carcinoma (IMC), or invasive cancer. **(C)** Kaplan–Meier overall survival curves for *in-vivo*-R *Arid1a* WT (N = 81), Het (N = 89; hazard ratio [HR] 1.9, 95% CI 1.3–2.8), and KO (N = 80; HR 2.3, 95% CI 1.5–3.5) mice. The curves were compared by log-rank test. **(D)** Representative H&E (top), ARID1A immunohistochemistry (middle), and YFP immunohistochemistry (bottom) of gastric tumors collected at time of death from *in-vivo*-R *Arid1a* WT, Het, and KO mice. SM indicates submucosa, M denotes muscularis externa. **(E)** Top: Representative gross photographs of stomachs from *in-vivo*-R *Arid1a* WT and KO mice illustrating the absence or presence, respectively, of extragastric extension (EGE) of gastric cancer through the outer serosal layer. Bottom: Proportion of *in-vivo*-R *Arid1a* WT, Het, and KO mice with EGE at time of death. Proportions were compared by Fisher’s exact test. **(F)** Top: Distribution of metastatic sites (lymph nodes [LN], liver, and lungs) by genotype at time of death. Bottom: Proportion of *in-vivo*-R *Arid1a* WT, Het, and KO mice with gross metastases at time of death. Proportions were compared by Fisher’s exact test. **(G)** Representative gross photographs (top), H&E (middle), and ARID1A immunohistochemistry (bottom) of liver and lung metastases from *in-vivo*-R mice. Yellow arrowheads indicate metastatic nodules. Yellow stars mark metastatic carcinoma; green crosses indicate adjacent normal tissue. **(H)** Success rate of generating primary cell lines from gastric tumors of *in-vivo-*R *Arid1a* WT, Het, and KO mice. **(I)** Representative genotyping PCR for *Arid1a* in *in-vivo-*R *Arid1a* Het and KO cell lines, showing floxed, wild-type, and deleted bands. +: positive control using non-recombined *Arid1a* Het tissue. **(J)** Western blot for ARID1A, E-CADHERIN, TRP53, and β-ACTIN in *in-vivo-*R *Arid1a* Het and KO cell lines. See also Supplemental Figure 2. **(K)** Representative brightfield (top) and immunofluorescence (IF; bottom) images of *in-vivo-*R *Arid1a* Het and KO cell lines. **(L)** Proportion of *in-vivo-*R *Arid1a* KO cell clones with successful *ARID1A* overexpression. **(M)** Western blot for ARID1A and β-ACTIN in *in-vivo-*R *Arid1a* KO cells with *ARID1A* overexpression (OE). +: positive control from *ARID1A*-expressing cell line. **(N)** Representative genotyping PCR for *Arid1a* in *in-vivo-*R *Arid1a* WT and KO tumoroids. **(O)** Western blot for ARID1A, E-CADHERIN, TRP53, and β-ACTIN in *in-vivo-*R *Arid1a* WT and KO tumoroids. **(P)** Representative brightfield (top) and IF (bottom) images of *in-vivo-*R *Arid1a* WT and KO tumoroids. **(Q)** Representative H&E (top) and ARID1A immunohistochemistry (bottom) in tumors generated from NSG mice injected with *in-vivo-*R *Arid1a* WT or KO tumoroids. See also Supplemental Figure 2. **(R)** Proportion of NSG mice that developed gross lung metastases following tail vein injection of *in-vivo-*R *Arid1a* WT or KO tumoroids. Proportions compared via Fisher’s exact test. See also Supplemental Figure 2. **(S)** Proportion of NSG mice that developed metastases following flank injection of *in-vivo-*R *Arid1a* KO cells overexpressing (OE) *ARID1A* or *GFP*. Proportions compared via Fisher’s exact test. **(T)** Proportion of NSG mice that developed metastases following flank injection of *in-vivo-*R *Arid1a* Het or KO cell lines. Proportions compared via Fisher’s exact test. **(U)** Tumoroid size over time measured by live imaging for *in-vivo-*R *Arid1a* WT and KO tumoroids. Data shown as mean ± SEM. Statistical comparison of pooled WT (N = 168) vs pooled KO (N = 276) genotypes by mixed-effects model (REML) with Geisser-Greenhouse correction. The time × genotype interaction was not significant. **(V)** Quantification of Ki-67^+^YFP^+^ cells as a percentage of total YFP^+^ cells in *in-vivo-*R *Arid1a* WT and KO tumoroids. Data shown as individual data points with mean ± SEM and compared with Welch’s t-test. **(W)** Quantification of Ki-67^+^ tumor cells as a percentage of total tumor cells in gastric cancers from *in-vivo-*R *Arid1a* WT, Het, and KO mice. Data shown as individual data points with mean ± SEM and compared with Kruskal-Wallis test. * P < 0.05, ** P < 0.01, **** P < 0.0001.

*In-vivo-*R *Arid1a* WT, Het, and KO mice all appeared grossly normal and were fertile. However, when we performed histologic examinations of their stomach mucosa, there was evidence of dysplasia in the *Arid1a* Het and *Arid1a* KO mice by day 5 of life (**Figure 2B**). By 6 weeks of age, some mice had intramucosal carcinoma in all three genotypes, and by 24 weeks, invasive cancer was present (**Figure 2B**). Although there was a trend toward accelerated cancer formation in *Arid1a* Het and *Arid1a* KO mice at various ages, this difference was not statistically significant as compared to the *Arid1a* WT cohort, likely due to small sample size.

To perform a statistically rigorous comparison, we generated a large cohort of *in-vivo-*R *Arid1a* WT, Het, and KO mice for long-term aging. As these mice aged, we observed that gastric cancer formed in all three genotypes with 100% penetrance; however, the mice demonstrated progressively shorter survival with the loss of each *Arid1a* allele. The median overall survival for *in-vivo*-R *Arid1a* WT mice was 42.6 weeks (N = 81), while the median survival of *Arid1a* Het mice was 39.4 weeks (N = 89, hazard ratio (HR), 1.9, 95% confidence interval (CI), 1.3–2.8), and of *Arid1a* KO mice was 38 weeks (N = 80, HR, 2.3, 95% CI, 1.5–3.5; P < 0.001; **Figure 2C**). The data presented are pooled from two cohorts of experimental mice generated at two separate time points and showed similar survival differences across the genotypes.

The tumors that formed were a mix of intestinal and diffuse histology, and the proportion of each type did not differ across the genotypes (**Figure 2D**). Using immunohistochemistry, we confirmed ARID1A protein expression was completely lost in the gastric cancers present in *in-vivo-*R *Arid1a* KO mice and retained in the majority of *Arid1a* WT and Het malignant cells. All tumors expressed YFP as expected (**Figure 2D**).

Consistent with previous work showing that tumor *Arid1a* loss promoted invasive capacity and metastasis,^21^ we found that the cancers that developed in the transgenic mice had increased invasiveness with *Arid1a* loss. At the time of death, only 11% of *in-vivo-*R *Arid1a* WT mice had extragastric extension of the tumor, which is analogous to stage T4 disease in human patients where the primary tumor erodes through the outermost serosal layer of the stomach wall, while 25% of *Arid1a* Het and 56% of *Arid1a* KO mice showed extragastric extension (P < 0.01; **Figure 2E**). Further supporting the concept that *Arid1a* loss promoted tumor invasiveness, 16% of *in-vivo-*R *Arid1a* WT had gross metastases at necropsy, as compared to 30% of *Arid1a* Het, and 60% of *Arid1a* KO mice (P < 0.01; **Figure 2F–G**). Metastases were seen in lymph nodes, peritoneum, liver, and lungs, which is a pattern of spread consistent with human disease.

To facilitate mechanistic work, we generated primary cell lines from *in-vivo-*R primary tumors. While we had a 50% success rate deriving cell lines from *Arid1a* Het and KO tumors, we could not generate any *Arid1a* WT cell lines, suggesting a strong selection pressure against *Arid1a* consistent with tumor suppressor effects (**Figure 2H–K; Supplemental Figure 2A**). We next tried overexpressing *ARID1A* in the *in-vivo-*R *Arid1a* KO cells. Of the 24 clones we generated, only 2 expressed ARID1A on Western blotting, again suggesting strong selection pressure against the presence of *Arid1a* (**Figure 2L–M; Supplemental Figure 2B**).

In contrast, we were successful in our attempts to generate three-dimensional tumoroids from both *in-vivo-*R *Arid1a* WT and KO tumors suggesting that the three dimensionality of the organoid system is more supportive of *in vitro* growth of *Arid1a* WT gastric cancer cells (**Figure 2N–P**). When we injected *in-vivo-*R *Arid1a* WT and KO tumoroids into the flanks of NSG mice, both genotypes formed tumors that had histologic features of gastric adenocarcinoma, had the expected ARID1A expression profile, and expressed YFP (**Figure 2Q; Supplemental Figure 2C**).

To confirm that these *in-vivo-*R tumoroids and cell lines maintained the relevant biological features seen in the transgenic mice, such as metastatic capability, we performed tail vein injections of tumoroids into NSG mice. We found that 13% of mice injected with *in-vivo-*R *Arid1a* WT tumoroids and 100% of mice injected with *Arid1a* KO tumoroids developed gross lung metastases (P < 0.0001; **Figure 2R; Supplemental Figure 2D**). Similarly, when we injected *in-vivo-*R *Arid1a* KO cell lines overexpressing *ARID1A* or *GFP* into the flanks of NSG mice, we found that only 13% of mice that were engrafted with *Arid1a* KO cells that overexpressed *ARID1A* developed metastases, as compared to 88% of mice engrafted with *Arid1a* KO cells expressing *GFP* (P < 0.05; **Figure 2S**). Finally, we saw a similar, but statistically insignificant, trend with *in-vivo-*R *Arid1a* Het and KO cell lines, which developed metastases at a rate of 40% and 76%, respectively (P = 0.07; **Figure 2T**), suggesting a dose-dependent effect of *Arid1a* as a metastasis suppressor. In sum, these data in the *in-vivo-*R context further confirmed that *Arid1a* acts as a tumor suppressor in gastric cancer, consistent with observations made in the *in-vitro-*R *Arid1a* tumoroids (**Figure 1E and 1H**).

In contrast to *in-vitro*-R tumoroids, when we compared the growth of *in-vivo-*R *Arid1a* WT and KO tumoroid lines *in vitro* via live imaging we did not observe a clear trend (**Figure 2U**). We then measured the Ki-67 positivity rates of *in-vivo-*R *Arid1a* WT and KO tumoroids injected into the flanks of NSG mice and did not observe a difference (**Figure 2V**), as was the case when we measured Ki-67 rates in the primary tumors of *in-vivo-*R mice (**Figure 2W**). Previous published work similarly led to contradictory conclusions regarding the effect of tumor *Arid1a* loss on proliferative capacity.^5,7,22^ These conflicting results, along with our findings, suggest that there are context-dependent factors driving the cell-autonomous tumor proliferative capacity in response to *Arid1a* loss.

### *In vivo* tumor *Arid1a* loss conferred immune evasive capability

To identify the molecular programs that underpin the aggressive phenotypes seen with *Arid1a* loss in mice, we performed bulk RNA-sequencing of the primary tumors collected from mice at the time of death. We performed gene set enrichment analysis (GSEA)^23^ and found that the most enriched programs in *in-vivo-*R *Arid1a* WT mouse tumors relative to *in-vivo-*R *Arid1a* KO tumors were immune-related (**Figure 3A**), suggesting an interaction between tumor *Arid1a* status and the host immune system that determined cancer behavior. Notably, we found that the HALLMARK_INTERFERON_GAMMA_RESPONSE pathway was the most enriched gene set in the murine *in-vivo-*R *Arid1a* WT tumors (**Figure 3A–B**). As IFN-γ is a key promoter of immune clearance of cancer through cytotoxic T cell activity,^24,25^ we performed qPCR to measure genes related to cytotoxic T cell activity. *Cxcl9*, *Cxcl10*, *Gzma*, and *Nkg7* were each expressed at higher levels in *Arid1a* WT tumors than KO tumors (**Figure 3C**).

**Figure 3.**
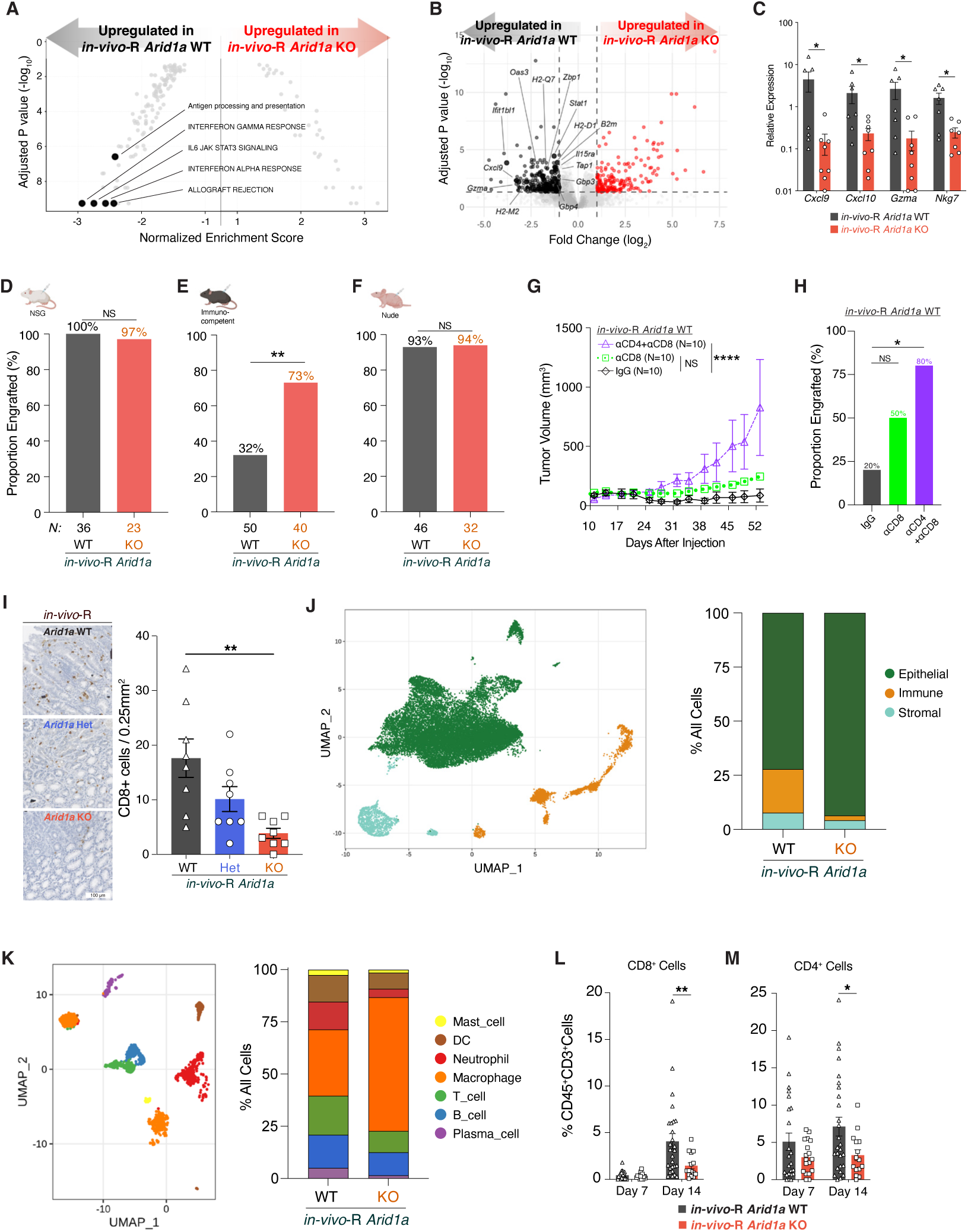
Tumor *Arid1a* loss promoted T cell–dependent immune evasion and an immune desert microenvironment. **(A)** Gene set enrichment analysis of bulk RNA-sequencing data from primary gastric tumors collected from *in-vivo-*R *Arid1a* WT and KO mice at time of death. Significantly enriched immune-related gene sets are highlighted. **(B)** Volcano plot showing differentially expressed genes between *in-vivo-*R *Arid1a* WT and KO primary tumors with genes in the INTERFERON_GAMMA_RESPONSE hallmark highlighted. **(C)** qPCR of selected genes. Data shown as mean ± SEM and compared with Mann-Whitney U test. **(D-F)** *in-vivo-*R *Arid1a* WT or KO tumoroids injected into the flanks of **(D)** NSG **(E)** syngeneic immunocompetent, and **(F)** athymic nude mice. Proportions of mice that developed tumors were compared via Fisher’s exact test. **(G)** Tumor growth curves following flank injection of *in-vivo*-R *Arid1a* WT tumoroids into immunocompetent mice treated with control IgG, anti-CD8 antibody, or combined anti-CD4 and anti-CD8 antibodies. Data shown as mean ± SEM. Statistical comparison versus IgG control through day 52 by mixed-effects model (REML) with Geisser-Greenhouse correction; asterisks denote the time × treatment interaction. **(H)** Proportion of immunocompetent mice that developed engrafted tumors following flank injection of *in-vivo-*R *Arid1a* WT tumoroids after treatment. Proportions compared via Fisher’s exact test. **(I)** Representative immunohistochemistry (left) and quantification (right) of CD8^+^ T cells in stomach sections from 24-week-old *in-vivo-*R *Arid1a* WT, Het, and KO mice. Data shown as mean ± SEM. Kruskal-Wallis test (P < 0.01) with Dunn’s multiple comparisons test (WT vs Het, P = non-significant, and WT vs KO, P < 0.01). **(J)** Left: UMAP visualization of single-cell RNA-sequencing data from 22,836 cells from 4 *in-vivo-*R *Arid1a* WT and 3 KO primary tumors. Right: Proportional representation of epithelial, immune, and stromal cell populations. **(K)** Left: UMAP visualization of single-cell RNA-sequencing data from 2,265 immune cells from 4 *in-vivo-*R *Arid1a* WT and 3 KO primary tumors. Right: Proportional representation of immune cell subtypes. DC: dendritic cells. **(L-M)** Flow cytometric analysis of tumors from *in-vivo-*R *Arid1a* WT and KO tumoroids 7 and 14 days after flank injection into syngeneic immunocompetent mice, measuring for infiltrated **(L)** CD8^+^ cells, **(M)** CD4^+^ cells. Data shown as mean ± SEM and compared using Welch’s t-test at each timepoint. NS, non-significant, * P < 0.05, ** P < 0.01, **** P < 0.0001.

We next tested the capacity of *in-vivo*-R tumoroids for immune evasion based on *Arid1a* status. First, we injected *in-vivo-*R *Arid1a* WT and KO tumoroids into the flanks of NSG mice and found that both genotypes engrafted at nearly a 100% rate (**Figure 3D**). In contrast, when we injected the tumoroids into the flanks of immunocompetent, syngeneic hosts, only 32% of *in-vivo-*R *Arid1a* WT tumoroids engrafted while 73% of *in-vivo-*R *Arid1a* KO tumoroids formed tumors (P < 0.01; **Figure 3E**). We confirmed these findings using the cell line systems, which were also generated from the primary tumors of transgenic mice. We found that 94% of NSG mice injected with *in-vivo-*R *Arid1a* KO cells overexpressing *GFP* or *ARID1A* formed tumors (**Supplemental Figure 3A**). In contrast, only 38% of immunocompetent mice injected with *ARID1A*-overexpressing *in-vivo-*R *Arid1a* KO cells formed tumors versus 75% of immunocompetent mice injected with control *GFP*-overexpressing cells (P < 0.05; **Supplemental Figure 3B**). Similarly, we found that 100% of NSG mice injected with *in-vivo-*R *Arid1a* Het and 97% of NSG mice injected with *in-vivo-*R *Arid1a* KO cell lines formed tumors (**Supplemental Figure 3C**). In contrast, when injected into immunocompetent hosts, only 27% of mice with *in-vivo*-R *Arid1a* Het cells and 96% of mice injected with *in-vivo-*R *Arid1a* KO cells engrafted (P < 0.0001; **Supplemental Figure 3D**).

Next, we tested if the immune system also affected *Arid1a*-mediated metastasis. We previously found that *in-vivo-*R *Arid1a* WT tumoroids that were injected into the tail veins of NSG mice had significantly fewer lung metastases as compared to mice injected with *in-vivo*-R *Arid1a* KO tumoroids (**Figure 2R**). When we performed tail vein injections of *in-vivo-*R *Arid1a* WT and KO tumoroids into immunocompetent recipients, we found no lung metastases in either genotype suggesting that the autochthonous derived tumoroids could not colonize the lungs in the setting of an intact immune system (data not shown). Thus, we utilized a different metastasis model in which tumor cells were introduced via intracardiac injections. First, we injected *in-vivo-*R *Arid1a* KO cells overexpressing *GFP* or *ARID1A* into NSG mice. At the time of sacrifice, 38% of mice bearing *ARID1A* overexpressing cells had metastasis, as compared to 79% of mice injected with *GFP*-expressing cells (P = 0.05; **Supplemental Figure 3E**), consistent with our earlier finding that *ARID1A* is a metastasis suppressor in the tail vein injection system (**Figure 2R**). However, when we used immunocompetent hosts as recipients, none of the mice bearing *ARID1A* overexpressing cells had metastasis, while 75% of mice injected with *GFP*-expressing cells formed metastases (P < 0.001; **Supplemental Figure 3E**). Notably, while a similar proportion of NSG and immunocompetent hosts developed metastases when injected with *GFP*-expressing cells, a significantly lower proportion of immunocompetent mice injected with *ARID1A*-expressing cells developed metastases, as compared to NSG hosts (38% vs 0%; P < 0.05; **Supplemental Figure 3E**). We confirmed this finding using *in-vivo-*R *Arid1a* Het and KO cell lines. While 75% of both NSG and immunocompetent mice developed metastases after injection with *in-vivo-*R *Arid1a* KO cells, the proportion of metastasis development significantly decreased with the injection of *in-vivo-*R *Het* cells from 57% in NSG mice to 6% in immunocompetent mice (P < 0.05; **Supplemental Figure 3F**). In sum, these data demonstrate that tumor cells in which *Arid1a* was lost in the presence of immune selection pressure (as accomplished by *in vivo* recombination of floxed *Arid1a* alleles), evaded immune clearance more effectively than *Arid1a* WT cells.

### *Arid1a-*mediated immune evasion required T cells

Since T cells are key effectors of the cancer immunity cycle, we next asked if T cells were necessary for *Arid1a*-mediated immune surveillance. We injected tumoroids and cell lines into athymic nude mice, which lack mature T cells. We found that *in-vivo*-R *Arid1a* WT and KO tumoroid injections had 93% and 94% engraftment rates, respectively (**Figure 3F**), while *in-vivo-* R *Arid1a* KO cells overexpressing *GFP* and *ARID1A* engrafted at a rate of 100% and 83%, respectively (**Supplemental Figure 3G**), and *in-vivo*-R *Arid1a* Het and KO cell lines engrafted at a rate of 94% and 100%, respectively (**Supplemental Figure 3H**; P = non-significant in all 3 experiments).

To further confirm the essential role of T cells in *Arid1a*-mediated immune clearance, we injected *in-vivo-*R *Arid1a* WT and KO tumoroids into immunocompetent hosts that were treated with anti-CD4 and anti-CD8 antibodies (**Supplemental Figure 3I**). We found that while mice injected with anti-CD8 antibody showed modestly increased *in-vivo-*R *Arid1a* WT tumoroid growth and engraftment (P = NS), concomitant depletion of CD4 and CD8 cells markedly increased tumor growth (P < 0.0001; **Figure 3G**) and engraftment (P < 0.05; **Figure 3H**). Of note, we found that while anti-CD4 and anti-CD8 treatment did increase *in-vivo-*R *Arid1a* KO tumoroid engraftment and tumor growth, it was to a lesser degree since *in-vivo-*R *Arid1a* KO tumoroids already robustly engrafted and grew in immunocompetent hosts (**Supplemental Figure 3J–3K**). In sum, these data showed that T cells were necessary for immune clearance of tumor cells that retained *Arid1a* expression.

Overall, the disparate immune phenotypes between the *in-vitro-*R and *in-vivo-*R systems suggest that the functional effects of tumor *Arid1a* on cancer immunity are dependent on the context in which *Arid1a* deletion occurred.

### *In vivo* tumor *Arid1a* loss promoted an immune desert microenvironment

Having confirmed that tumor *Arid1a* status played an essential role in immune surveillance of gastric cancer, we sought to delineate the effects of *in vivo* tumor *Arid1a* loss on the tumor immune microenvironment. As we already found that T cells were necessary in *Arid1a*-mediated immune evasion, we measured the number of CD8^+^ T cells in the stomachs of transgenic mice. Rather than examining mice at the time of death, which was highly variable within and across genotypes, we examined the stomachs of 24-week-old mice, which is an age that we previously showed that these mice have invasive cancers on histology but not grossly (**Figure 2B**). We found that there was a progressive decrease in the number of CD8^+^ T cells in the stomachs of *Arid1a* WT, Het, and KO mice, respectively (P < 0.01; **Figure 3I**). We compared the immune microenvironments of 4 *in-vivo-*R *Arid1a* WT and 3 KO primary tumors via single-cell RNA sequencing (scRNA-seq) of a total of 22,836 cells. Immune cells comprised 20% of the total population in *Arid1a* WT tumors, compared to only 2% in *Arid1a* KO tumors (**Figure 3J**). These data, in sum, suggest that tumor *Arid1a* loss induced an immune desert phenotype.^26^

We characterized the immune cell population in more detail and found that in *in-vivo-*R *Arid1a* KO tumors, there was an expansion of the macrophage population and a decrease in the proportion of dendritic cells, T cells, and neutrophils (**Figure 3K**). Next, we used flow cytometry to analyze the immune cell infiltration induced by *in-vivo*-R *Arid1a* WT and KO tumoroids injected into the flanks of immunocompetent hosts 7 and 14 days post-injection. Of note, even though most *in-vivo-*R *Arid1a* WT injections eventually regress (**Figure 3E**), this immune elimination phase takes place over a 3- to 4-week time frame and thus at 7 and 14 days post-injection, the tumors were still present. While we did not see a difference in CD45^+^ immune cell infiltration between *in-vivo*-R *Arid1a* WT and KO tumoroids at either time point (**Supplemental Figure 3L**), we did note a difference in T cell infiltration consistent with the scRNA-seq findings. At day 7, there were very few CD8^+^ cells in both *in-vivo*-R *Arid1a* WT and KO tumors, but by day 14, there were significantly more CD8^+^ cells in the *in-vivo*-R *Arid1a* WT tumors (P < 0.01; **Figure 3L**). Similarly, there was no difference in the number of CD4^+^ cells at day 7, but by day 14, there were significantly more CD4^+^ cells in the *in-vivo*-R *Arid1a* WT tumors (P < 0.05; **Figure 3M**).

These results are consistent with our earlier findings demonstrating that T cells are key mediators in *Arid1a* induced immunity (**Figure 3E–H**) and align with the observed difference in CD8^+^ T-cell infiltration in transgenic mice stomachs (**Figure 3I**).

Within the myeloid compartment, while we did not find a difference in pro-inflammatory F4/80⁺CD11b⁺CD206⁻ macrophages (**Supplemental Figure 3M**), we did observe that *in-vivo*-R *Arid1a* KO tumors had significantly more immunosuppressive F4/80⁺CD11b⁺CD206^+^ macrophages 7 days post-injection than *Arid1a* WT tumors (P < 0.05; **Supplemental Figure** 3N).^27,28^

### Tumor *Arid1a* loss blunted GM-CSF secretion and type II interferon response

Since ARID1A is an essential component of the SWI/SNF chromatin remodeling complex, we next performed ATAC-seq to compare the chromatin accessibility of *in-vivo*-R *Arid1a* WT and KO tumoroids to identify the molecular mechanisms that are responsible for *Arid1a*-mediated immune evasion. Out of 81,751 consensus peaks identified, 23,758 were significantly different between *in-vivo*-R *Arid1a* WT and KO tumoroids. Only 464 peaks were upregulated, while 23,294 peaks were downregulated in the *in-vivo*-R *Arid1a* KO lines. Thus, *Arid1a* loss induced an overall more closed chromatin state in our gastric cancer tumoroids (**Figure 4A**), a finding that is consistent with previous work done by us and others that show *Arid1a* loss reduces chromatin accessibility.^21,22^

**Figure 4.**
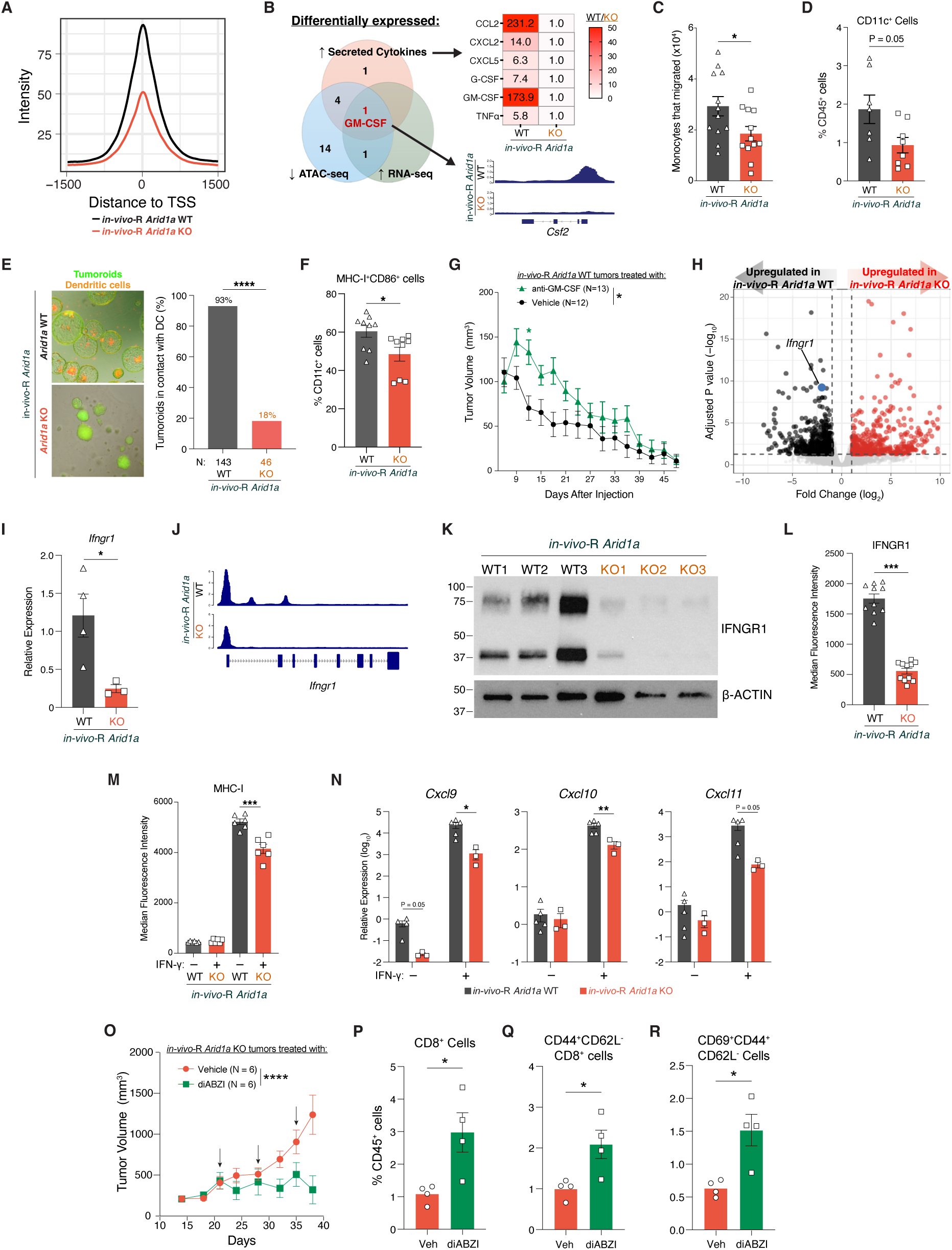
Tumor *Arid1a* induced GM-CSF secretion and type II interferon response to spur the cancer immunity cycle. **(A)** Assay for transposase-accessible chromatin using sequencing (ATAC-seq) analysis showing chromatin accessibility profiles relative to transcription start sites (TSS) in *in-vivo-*R *Arid1a* WT and KO tumoroids. **(B)** Integrated analysis of differential cytokine gene expression, secretion, and chromatin accessibility in *in-vivo-*R *Arid1a* WT and KO tumoroids. Venn diagram shows cytokines with >5-fold differential secretion, significant differential expression by RNA-seq, and differential accessibility by ATAC-seq. Table shows fold-change in secretion for the top differentially secreted cytokines as a ratio of WT/KO. GM-CSF is the only cytokine with concordant changes across all three modalities. The track for *Csf2* (which encodes GM-CSF) is shown. **(C)** Monocyte migration toward conditioned media from *in-vivo-*R *Arid1a* WT or KO tumoroids. Data shown as mean ± SEM and compared with Welch’s t-test. **(D)** Flow cytometric quantification of CD11c^+^ dendritic cells in tumor-draining lymph nodes of mice bearing *in-vivo-*R *Arid1a* WT or KO tumoroids. Data shown as mean ± SEM and compared with Welch’s t-test. **(E)** Representative images and quantification of dendritic cells in direct contact with *in-vivo-*R *Arid1a* WT or KO tumoroids during co-culture. Proportions compared via Fisher’s exact test. **(F)** Flow cytometric analysis of dendritic cell activation markers (MHC-I^+^CD86^+^) following co-culture with *in-vivo-*R *Arid1a* WT or KO tumoroids. Data shown as mean ± SEM and compared with Welch’s t-test. **(G)** Tumor growth curves following flank injection of *in-vivo*-R *Arid1a* WT tumoroids into immunocompetent mice treated with vehicle or anti–GM-CSF antibody. Data shown as mean ± SEM. Curves compared by mixed-effects model (REML) with Geisser-Greenhouse correction. The black asterisk denotes a significant time × treatment interaction. Individual timepoints were compared with Bonferroni’s multiple comparisons test and day 12 was the only timepoint that was significantly different between groups (green asterisk). **(H)** Volcano plot showing differential gene expression between *in-vivo-*R *Arid1a* WT and KO tumoroids from bulk RNA-seq. **(I)** qPCR analysis of *Ifngr1* expression in *in-vivo-*R *Arid1a* WT and KO tumoroids. Data shown as mean ± SEM and compared with Welch’s t-test. **(J)** ATAC-seq tracks at the *Ifngr1* locus in *in-vivo-*R *Arid1a* WT and KO tumoroids. **(K)** Western blot for IFNGR1 and β-ACTIN in *in-vivo-*R *Arid1a* WT and KO tumoroids. **(L)** Flow cytometric analysis of cell-surface IFNGR1 expression on *in-vivo-*R *Arid1a* WT and KO tumoroids. Left: representative histogram. Right: quantification of median fluorescence intensity. Data shown as mean ± SEM and compared with Welch’s t-test. **(M)** Flow cytometric analysis of cell-surface MHC-I expression on *in-vivo-*R *Arid1a* WT and KO tumoroids at baseline and following IFN-γ stimulation. Left: representative histogram. Right: quantification of median fluorescence intensity. Data shown as mean ± SEM and compared with Welch’s t-test. **(N)** qPCR analysis of select genes following IFN-γ treatment of *in-vivo-*R *Arid1a* WT and KO tumoroids. Data shown as mean ± SEM and compared with Welch’s t-test. **(O)** Tumor growth curves following flank injection of *in-vivo*-R *Arid1a* KO tumoroids in mice treated with vehicle or diABZI. Treatment (arrows) was administered at days 21, 28, and 35 after tumoroid injection. Data shown as mean ± SEM. Statistical comparison of vehicle vs diABZI from day 21 through day 38 by mixed-effects model (REML) with Geisser-Greenhouse correction. Asterisks denote the time × treatment interaction. **(P-R)** Flow cytometric analysis of *in-vivo*-R *Arid1a* KO tumors following intratumoral diABZI treatment, quantifying as a percentage of CD45+ cells with data shown as mean ± SEM and compared with Welch’s t-test. **(P)** CD8+ T cells, **(Q)** CD44+CD62L– effector memory CD8+ T cells, and **(R)** CD69+CD44+CD62L– activated effector memory T cells. * P < 0.05, ** P < 0.01, *** P < 0.001, **** P < 0.0001.

A key step to initiate the cancer immunity cycle is the recruitment of immune cells to the tumor immune microenvironment to unleash a feed forward cycle for further activation and recruitment of anti-tumor actors, a process that is supported by cytokine secretion by cancer and immune cells.^29^ When we examined the loci of 32 key cytokines, we found that 20 were significantly closed with *Arid1a* loss (**Figure 4B**). Next, we measured cytokine levels in conditioned media from *in-vivo-*R *Arid1a* WT and KO tumoroids and we found markedly different inflammatory secretomes, with *in-vivo-*R *Arid1a* WT tumoroids producing 6 cytokines that were at least 5-fold higher than *Arid1a* KO tumoroids. Of these 6 cytokines, 5 had more closed loci in *in-vivo-*R *Arid1a* KO tumoroids on ATAC-seq (**Figure 4B**). Finally, when we analyzed bulk RNA-seq data from the tumoroids, we found that of these 5 cytokines with markedly closed chromatin in *Arid1a* KO tumoroids, only the expression of *Csf2*, which encodes GM-CSF, was significantly lower in *in-vivo-*R *Arid1a* KO tumoroids. This suggests that GM-CSF expression was most likely under the direct control of *Arid1a*, whereas the other differentially secreted cytokines may be due to transcriptional or post-translational modifications. Notably, GM-CSF was the second highest differentially secreted cytokine with *in-vivo-*R *Arid1a* WT tumoroids producing a 174-fold higher amount than *Arid1a* KO tumoroids (P < 0.05; **Figure 4B**).

GM-CSF plays a key role in dendritic cell recruitment and maturation.^30^ Thus, we asked if *in-vivo-*R *Arid1a* WT tumoroids can more effectively attract dendritic cells. We performed a migration assay for monocytes, which are one precursor of dendritic cells, using conditioned media produced by *in-vivo-*R *Arid1a* WT and KO tumoroids and found that media from *Arid1a* WT tumoroids led to significantly more monocyte migration (P < 0.05; **Figure 4C**). Next, we compared the number of dendritic cells in tumor draining lymph nodes of mice injected with *in-vivo-*R *Arid1a* WT and KO tumoroids and found that there were significantly more CD11c^+^ cells in the tumor draining lymph nodes of mice bearing *in-vivo-*R *Arid1a* WT tumoroids (P = 0.05; **Figure 4D**). We then co-cultured tumoroids directly with dendritic cells and found that 93% of *in-vivo-*R *Arid1a* WT tumoroids had dendritic cells touching them as compared to only 18% of *Arid1a* KO tumoroids (P < 0.0001; **Figure 4E**). Notably, we also found that a higher proportion of dendritic cells co-cultured with *Arid1a* WT tumoroids expressed activation markers than those co-cultured with *Arid1a* KO tumoroids (P < 0.05; **Figure 4F**). In sum, our findings suggest that tumor *Arid1a* loss reduced tumor GM-CSF secretion, which lessened dendritic cell recruitment into the tumor immune microenvironment, and blunted the initiation of the cancer immunity cycle to promote immune evasion.

Finally, we tested the necessity of GM-CSF for immune clearance of *Arid1a* WT cancer cells. We injected *in-vivo-*R *Arid1a* WT tumoroids into the flanks of immunocompetent host mice treated with anti-GM-CSF antibodies and found that while anti-GM-CSF treatment induced an initial growth of tumor in the first two weeks after injection, the tumors all eventually regressed (**Figure 4G**). These data demonstrated that other mechanisms must also be involved in *Arid1a*-mediated immune clearance as blocking GM-CSF alone was not sufficient to phenocopy *Arid1a* loss.

To identify other tumor cell-autonomous factors that contribute to *Arid1a*-mediated immune clearance beyond GM-CSF, we compared the RNA-seq data of *in-vivo-*R *Arid1a* WT and KO tumoroids and found that *Ifngr1* expression was significantly higher in *Arid1a* WT tumoroids (**Figure 4H**), which we validated with qPCR (**Figure 4I**). This finding correlated with the bulk RNA-seq data of the primary tumors from transgenic mice that showed the IFN-γ response pathway (HALLMARK_INTERFERON_GAMMA_RESPONSE; **Figure 3A**) was the most differentially expressed gene set between murine *Arid1a* WT and KO tumors. On review of the ATAC-seq data we found that the *Ifngr1* locus was more closed in the *Arid1a* KO tumoroids (**Figure 4J**). Finally, we confirmed that *Arid1a* KO tumoroids had reduced total IFNGR1 protein expression via Western blotting (**Figure 4K**) and less IFNGR1 on cell surface via flow cytometry (P < 0.001; **Figure 4L; Supplemental Figure 4A**).

IFNGR1 is the ligand binding component of the IFN-γ receptor heterodimer complex and is essential to initiating the type II interferon response.^31^ Thus, we sought to further interrogate the role that type II interferon signaling played in *Arid1a* mediated immune evasion. Based on the reduced expression of *Ifngr1*, we hypothesized that *in-vivo-*R *Arid1a* KO tumoroids would mount a blunted response to IFN-γ. We exposed *in-vivo-*R *Arid1a* WT and KO tumoroids to exogenous IFN-γ *in vitro* and measured cell surface MHC-I expression, which is upregulated as part of the type II interferon signaling response and a critical component of the tumor-immune escape mechanism. We found that while at baseline, there was very low MHC-I expression on the surface of both *Arid1a* WT and KO tumoroids, *Arid1a* WT tumoroids expressed significantly more cell surface MHC-I upon IFN-γ exposure (P < 0.001; **Figure 4M; Supplemental Figure 4B**). We also observed that IFN-γ treatment induced a significantly higher increase in *Cxcl9*, *Cxcl10*, and *Cxcl11* expression in *in-vivo-*R *Arid1a* WT tumoroids (**Figure 4N**), further confirming that *in-vivo-*R *Arid1a* KO tumoroids had blunted type II interferon response.

### *Arid1a*-null cancers can be eradicated with the induction of a robust immune response

We next sought to identify strategies to treat *Arid1a*-deficient tumors that evaded immune clearance by blunting the cancer immunity cycle initiation. Since we found that *Arid1a* loss led to tumor cell-autonomous reduction in GM-CSF secretion and *Ifngr1* expression, we expected that tumor *Arid1a* loss hinders immune clearance via local, rather than systemic, mechanisms. To test if *in-vivo-*R *Arid1a* WT tumoroids released or induced systemic factors that are sufficient to clear *Arid1a* KO tumoroids, we performed concomitant injection of both *in-vivo-*R *Arid1a* WT and KO tumoroids into contralateral flanks of the same mice, positing that if there were any systemic anti-tumor factors induced by *Arid1a* WT tumoroids, then the KO cells would also be eradicated. However, we found that 100% of *Arid1a* KO tumoroids engrafted, consistent with tumor *Arid1a* mediating only local effects (**Supplemental Figure 4C**).

Next, we tested PD-1/PD-L1 blockade as a therapeutic strategy against *Arid1a*-null cancer cells. Since we previously found that *in-vivo-*R *Arid1a* KO cancer cells excluded T cell infiltration into the tumors, we hypothesized that anti-PD-L1/PD-1 therapy will not affect tumor growth since this class of treatment requires the presence of T cells for effect. We first injected *in-vivo-*R *Arid1a* KO cells into the flanks of syngeneic immunocompetent mice, and upon the tumors reaching 200 mm^3^, the mice were treated with either control IgG or anti-PD-1 antibodies. We found no difference in tumor growth (**Supplemental Figure 4D**). Next, we treated *in-vivo-*R *Arid1a* WT and KO mice with PBS or anti-PD-L1 antibody. We initiated weekly injections at 24 weeks of age, when these mice were known to have at least intramucosal cancer (**Figure 2B**) and again, found no difference in survival **(Supplemental Figure 4E**).

Since *Arid1a* mediated local effects in the tumor immune microenvironment, we hypothesized that provoking a robust local immune reaction would eradicate *in-vivo-*R *Arid1a* KO cells. To test this theory, we leveraged the adaptive immune system that establishes immunologic memory which provides a rapid and robust response upon re-exposure to antigens. We injected *in-vivo-* R *Arid1a* WT tumoroids into immunocompetent mice and waited 80 days, during which time immunologic memory could develop. Using mice that cleared this initial exposure, we re-challenged them with *in-vivo-*R *Arid1a* KO tumoroids injected into the contralateral flank. We found that only 38% of injections led to engraftment (**Supplemental Figure 4F**), which is markedly less than the 73% engraftment rate when *in-vivo*-R *Arid1a* KO tumoroids were injected into naïve hosts (P < 0.01; **Figure 3E**). These results showed that *in-vivo-*R *Arid1a* WT and KO tumoroids share neoantigens that primed the immune system to clear a re-exposure and confirm that *in-vivo-*R *Arid1a* KO tumoroids can be immunologically eradicated, provided that a strong enough immune reaction is induced.

The cGAS–STING pathway is a central innate immune sensing axis that induces type I interferon and pro-inflammatory cytokine programs to promote dendritic cell activation and T cell priming, thereby enhancing anti-tumor immunity.^32^ We tested intratumoral injections of the STING agonist diamidobenzimidazole (diABZI), as a therapeutic option.^33^ First, we engrafted *in-vivo-*R *Arid1a* KO cells into the flanks of syngeneic immunocompetent hosts and one day later injected diABZI into the tumor site, followed by weekly injections. We found that diABZI treatment prevented tumor growth (P < 0.0001; **Supplemental Figure 4G**). Notably, even once we stopped diABZI therapy, we saw only a low rate of tumor regrowth, demonstrating that this strategy can completely eradicate cancer cells. We next tested diABZI treatment in established tumors. We engrafted *in-vivo-*R *Arid1a* KO cells into the flanks of syngeneic immunocompetent mice, and upon the tumors reaching a mean size of 400 mm^3^, we performed intratumoral injection of diABZI, which provided significant tumor control (P < 0.0001; **Figure 4O**), with nearly half the tumors showing complete regression (**Supplemental Figure 4H**).

To confirm that diABZI initiated a robust immune response, we used flow cytometry to analyze immune cell infiltration after diABZI treatment. We found while there was no difference in viable immune cells (**Supplemental Figure 4I**), there was a trend toward a higher proportion of CD3^+^ T cells (P = 0.06; **Supplemental Figure 4J**), most notably in the CD8^+^ population (P < 0.05; **Figure 4P**). Importantly, diABZI treatment induced a higher proportion of T effector memory (T_EM_) cells (P < 0.05; **Figure 4Q**) and antigen experienced T_EM_ cells (P < 0.05; **Figure 4R**), which suggest an activated adaptive immunity response.

### *In vitro* tumor *Arid1a* deletion did not affect GM-CSF or type II interferon response

We next asked how *Arid1a* status in *in-vitro-*R tumoroids, which had *Arid1a* deleted *in vitro*, affected tumoroid cytokine production. We found that *in-vitro-*R *Arid1a* WT and KO tumoroids had a markedly different secretome pattern as compared to the *in-vivo-*R tumoroids. Most notably, GM-CSF production was not significantly different between the two genotypes in the *in-vitro-*R tumoroids (**Figure 5A**). Consistent with the lack of difference in GM-CSF secretion, we found that *in-vitro-*R *Arid1a* WT and KO tumoroids attracted a similar number of dendritic cells while in co-culture (66% vs 71%, respectively; P = NS; **Figure 5B**).

**Figure 5.**
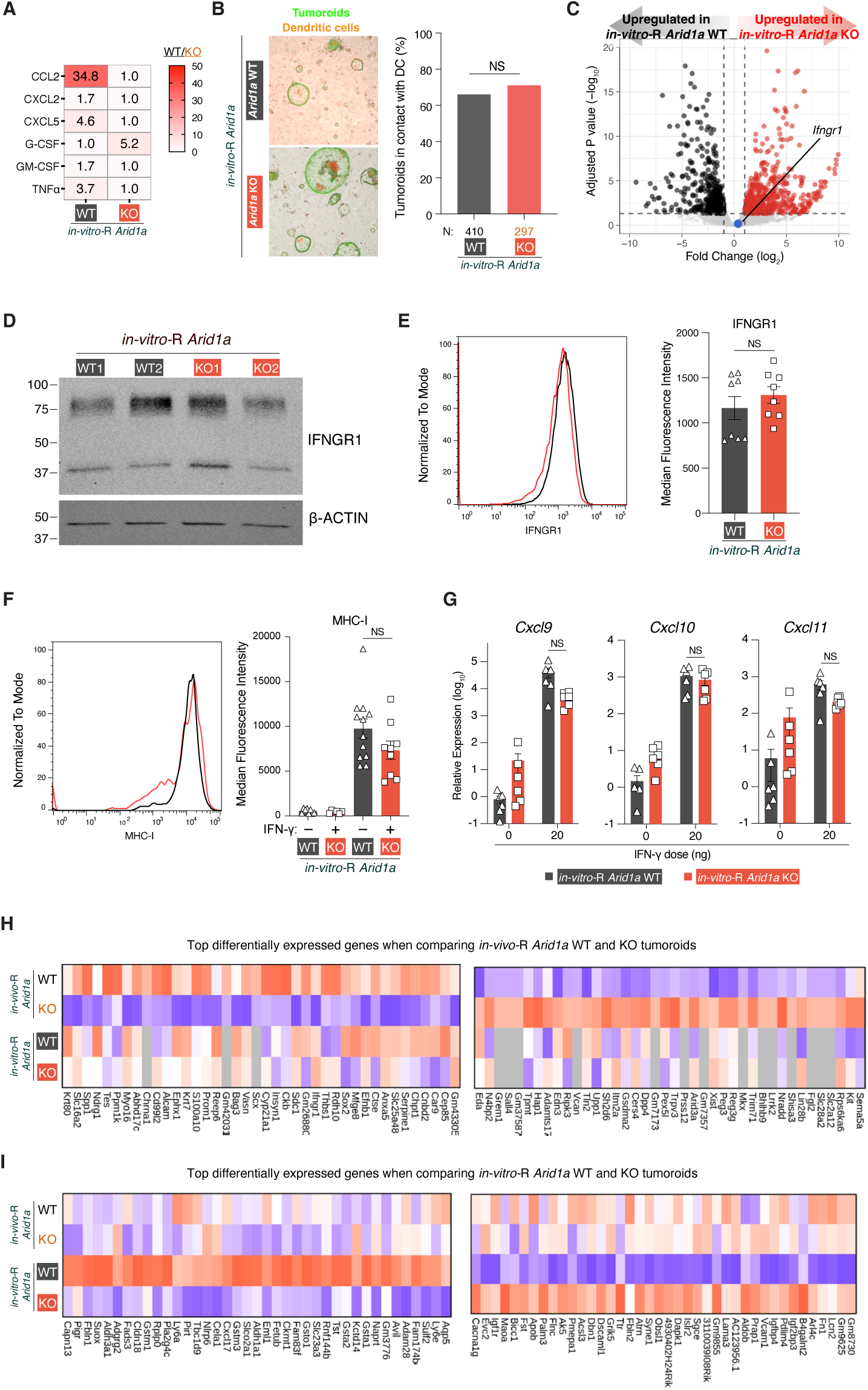
*In vitro Arid1a* deletion did not affect GM-CSF or type II interferon response. **(A)** Quantification of cytokines in conditioned media from *in-vitro-*R *Arid1a* WT and KO tumoroids. **(B)** Representative images and quantification of dendritic cells in direct contact with *in-vitro-*R *Arid1a* WT or KO tumoroids during co-culture. Proportions compared via Fisher’s exact test. **(C)** Volcano plot showing differential gene expression between *in-vitro-*R *Arid1a* WT and KO tumoroids from bulk RNA-seq. **(D)** Western blot for IFNGR1 and β-ACTIN in *in-vitro-*R *Arid1a* WT and KO tumoroids. **(E)** Flow cytometric analysis for cell-surface IFNGR1 expression on *in-vitro-*R *Arid1a* WT and KO tumoroids. Left: representative histogram. Right: quantification of median fluorescence intensity. Data shown as mean ± SEM and compared with Welch’s t-test. **(F)** Flow cytometric analysis for cell-surface MHC-I expression on *in-vitro-*R *Arid1a* WT and KO tumoroids at baseline and following IFN-γ stimulation. Left: representative histogram. Right: quantification of median fluorescence intensity. Data shown as mean ± SEM and compared with Welch’s t-test. **(G)** qPCR of select genes following IFN-γ treatment of *in-vitro-*R *Arid1a* WT and KO tumoroids. Data shown as mean ± SEM and compared with Welch’s t-test. **(H)** Heatmap showing the top differentially expressed genes between *in-vivo-*R *Arid1a* WT and KO tumoroids, plotted across both *in-vivo-*R *Arid1a* and *in-vitro-*R *Arid1a* tumoroid datasets. **(I)** Heatmap showing the top differentially expressed genes between *in-vitro-*R *Arid1a* WT and KO tumoroids, plotted across both *in-vivo-*R *Arid1a* and *in-vitro-*R *Arid1a* tumoroid datasets. NS, non-significant.

Next, we tested how *Arid1a* status in *in-vitro-*R tumoroids affected their response to IFN-γ exposure. First, RNA-seq of *in-vitro-*R *Arid1a* WT and KO tumoroids showed no difference in *Ifngr1* expression, which we confirmed with qPCR (**Figure 5C and Supplemental Figure 5A**). Consistent with the RNA level findings, we found that there was no clear difference in IFNGR1 protein expression between *in-vitro-*R *Arid1a* WT and KO tumoroids on Western blots (**Figure 5D**) or on flow cytometry (**Figure 5E**). Finally, when we exposed *in-vitro-*R tumoroids to exogenous IFN-γ, we did not find any differences between *Arid1a* WT and KO tumoroids in type II interferon responses as determined by cell surface MHC-I expression (**Figure 5F**) and induction of pro-inflammatory cytokines (**Figure 5G**).

We next compared the effects of *Arid1a* loss on the transcriptomes in *in-vivo-*R and *in-vitro-*R tumoroids. Using bulk RNA-seq we found that out of the top 80 most differentially expressed genes between *in-vivo-*R *Arid1a* WT and KO tumoroids, few showed similar differential expression patterns when *in-vitro-*R *Arid1a* WT and KO tumoroids were compared (**Figure 5H**).

Similarly, when we analyzed the top 80 most differentially expressed genes between *in-vitro-*R *Arid1a* WT and KO tumoroids, we did not see a similar expression pattern when *in-vivo-*R *Arid1a* WT and KO tumoroids were analyzed (**Figure 5I**). These data demonstrated that *Arid1a* loss induced different transcriptomic changes based on whether *Arid1a* deletion occurred *in vitro* versus *in vivo*.

### Tumor *ARID1A* mutation effect on human gastric cancer immunity differed based on molecular subtype

Human gastric adenocarcinoma is molecularly heterogeneous, with four major subtypes: Epstein–Barr virus-positive (EBV), microsatellite instability (MSI), chromosomal instability (CIN), and genomically stable (GS).^15^ EBV and MSI subtypes comprise 7-10% and 10-22%, respectively, of gastric adenocarcinoma, while the majority of gastric adenocarcinomas are CIN or GS.^15^ To determine which of the four TCGA subtypes our *in-vivo-*R *Arid1a* murine tumors most resembled, we developed a cross-species classification framework that projected mouse bulk RNA-seq expression profiles onto four TCGA groups (**Supplemental Figure 6A**). All 11 *in-vivo-*R *Arid1a* WT tumors were classified as GS with high predicted subtype probability. Among *Arid1a* Het samples, 6 of 7 (86%) were classified as GS and 1 of 7 (14%) as CIN. In the *Arid1a* KO group, 8 of 11 (73%) were classified as GS and 3 of 11 (27%) as CIN. No samples in any genotype were classified as EBV or MSI. Thus, we concluded that the *in-vivo-*R *Arid1a* tumors most resembled the GS subtype.

To determine the association between *ARID1A* mutations and cancer immunity in human patients, we interrogated The Cancer Genome Atlas gastric adenocarcinoma cohort (TCGA STAD) comprised of 440 samples, of which 50 were GS and 6 of the 50 (12%) harbored *ARID1A* loss-of-function (LOF) mutations. We performed GSEA comparing GS *ARID1A* WT and mutant tumors and found that mutants were depleted for immune related hallmark gene sets, such as ALLOGRAFT_REJECTION, INTERFERON_GAMMA_RESPONSE, and INTERFERON_ALPHA_RESPONSE, as compared to *ARID1A* WT samples (**Figure 6A; Supplemental Figure 6B**), consistent with the findings in our transgenic mouse model where tumor *Arid1a* loss resulted in immune-depleted phenotypes.

**Figure 6.**
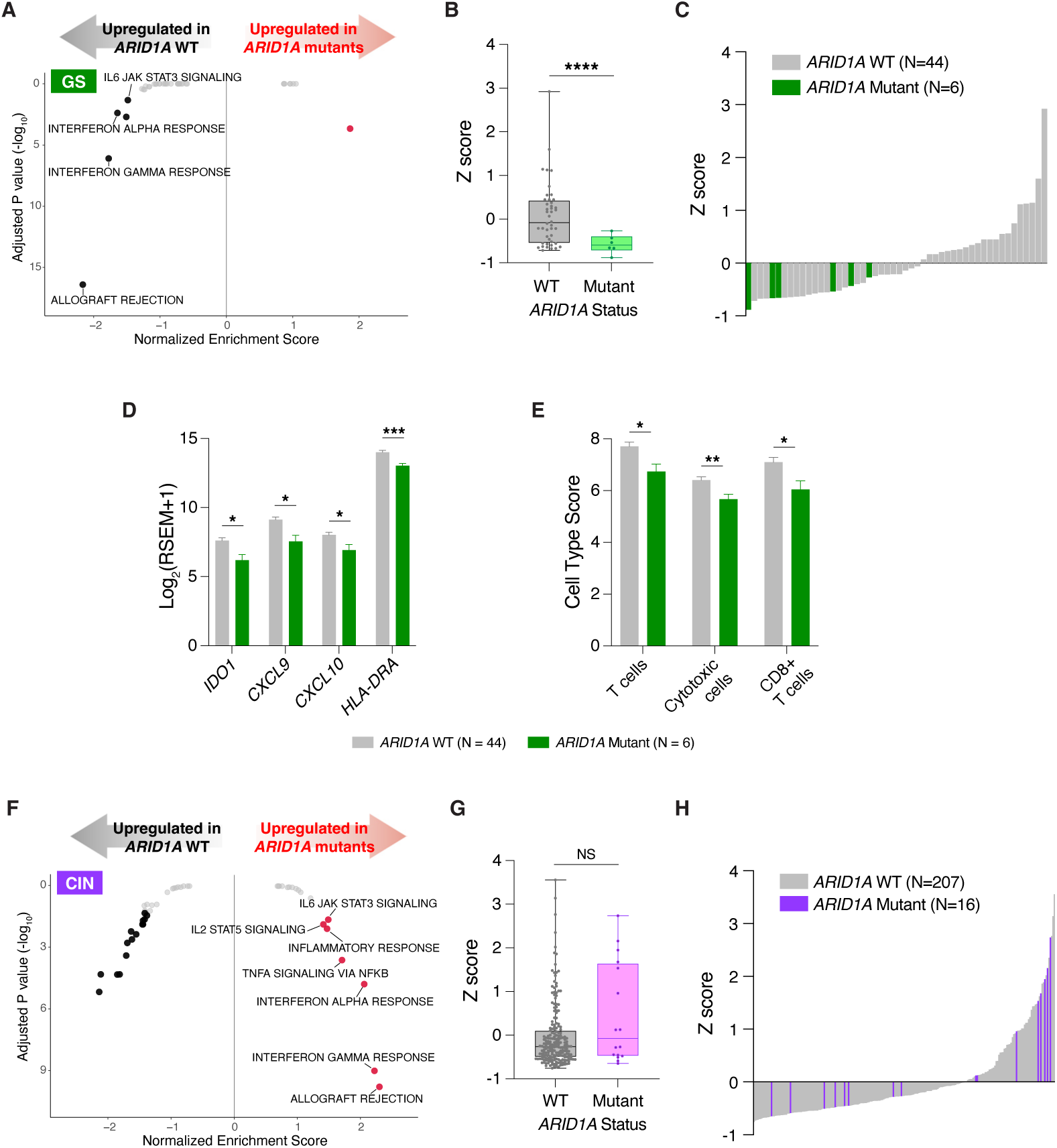
Effects of *ARID1A* loss-of-function mutations in human gastric cancer were dependent on molecular subtype. **(A)** Human gastric adenocarcinoma samples from The Cancer Genome Atlas were analyzed via gene set enrichment analysis (GSEA) comparing genomically stable (GS) tumors that carried *ARID1A* loss-of-function (LOF) mutations (N = 6) to *ARID1A* wild-type (WT) samples (N = 44). Select immune-related pathways are labeled. **(B)** 6-gene interferon-γ signature z-score comparison of GS *ARID1A* WT and mutant tumors. Data shown as a box-and-whisker plot (box, interquartile range; center line, median; whiskers, minimum to maximum) and compared with Welch’s t-test. **(C)** Waterfall plot of IFN-γ z-scores for all 50 GS tumors, ranked by score. *ARID1A* WT samples in gray; *ARID1A* mutant samples in green. **(D)** Expression of IFN-γ pathway genes in GS *ARID1A* WT and mutant tumors. Log-transformed data shown as mean ± SEM and compared with Welch’s t-test. **(E)** Immune cell type signature scores in GS *ARID1A* WT and mutant tumors. Data shown as mean ± SEM and compared with Welch’s t-test. **(F)** GSEA comparing chromosomal instability (CIN) tumors that carried *ARID1A* LOF mutations (N = 16) to *ARID1A* wild-type (WT) tumors (N = 207). Select immune-related pathways are labeled. **(G)** 6-gene interferon-γ signature z-score comparison of CIN *ARID1A* WT and mutant tumors. Data shown as a box-and-whisker plot (box, interquartile range; center line, median; whiskers, minimum to maximum) and compared with Welch’s t-test. **(H)** Waterfall plot of IFN-γ z-scores for all 223 CIN tumors, ranked by score. *ARID1A* WT samples in gray; *ARID1A* mutant samples in purple. NS, non-significant, * P < 0.05, ** P < 0.01, *** P < 0.001, **** P < 0.0001.

We next compared the level of IFN-γ–driven immune activation in the tumor microenvironment using a 6-gene IFN-γ signature and found that *ARID1A* LOF tumors had significantly lower IFN-γ signature scores compared to *ARID1A* WT tumors (mean −0.59 vs +0.06; P < 0.0001; **Figure 6B**).^34^ Notably, all 6 GS *ARID1A* LOF samples uniformly had negative IFN-γ signature z-scores (**Figure 6C**). When we examined the individual IFN-γ pathway genes we observed that *IDO1*, *CXCL9*, *CXCL10*, and *HLA-DRA* were all reduced in *ARID1A* mutant tumors as compared to WT tumors (all P < 0.05; **Figure 6D**). Finally, immune deconvolution analysis demonstrated significantly lower infiltration of T cells, cytotoxic cells, and CD8+ cells in *ARID1A* LOF mutant samples compared to *ARID1A* WT tumors (all P < 0.05; **Figure 6E**). Overall, these results were consistent with the immune exclusion phenotype observed in our *in-vivo-*R *Arid1a* KO model.

Next, we compared the immune phenotype of *ARID1A* mutants in the CIN subtype, which is the other major molecular subtype of gastric cancers. In direct contrast to the GS samples, GSEA comparing the 16 CIN *ARID1A* LOF mutants to the 207 CIN *ARID1A* WT samples showed that the *ARID1A* mutant samples were enriched for multiple immune pathways (**Figure 6F; Supplemental Figure 6C**) and there was no difference in the 6-gene IFN-γ signature scores (**Figure 6G**). Notably, when we reviewed the CIN-subtype *ARID1A* LOF tumors individually, we found that there were highly heterogeneous IFN-γ scores, with 8 samples scoring above and the other 8 scoring below zero (**Figure 6H**). Overall, these results were consistent with the concept that the immune effect of *ARID1A* mutations was related to the context in which they occur. In particular, the immunosuppressive phenotype associated with *ARID1A* LOF is specific to the GS molecular context, consistent with our findings in our mouse and tumoroid models.

## DISCUSSION

In this study, we compared two isogenic gastric cancer models that differed only in the context in which *ARID1A* was deleted and found markedly different effects on gastric cancer immunity. When tumor *Arid1a* loss occurred *in vitro*, *Arid1a* deletion did not affect the cancer cells’ immune evasion proficiency. However, in an autochthonous genetically engineered mouse model, in which tumor cells developed *in vivo* over months, *Arid1a*-null cancer cells capable of T cell-dependent immune escape emerged, leading to shortened survival for mice. Mechanistically, we found that autochthonous tumor *Arid1a* loss reprogrammed the immune microenvironment into a desert phenotype via blunting of GM-CSF production and type II interferon response. Finally, we confirmed that the correlation between *ARID1A* LOF mutations and immune activation in human gastric cancer samples was dependent on tumor molecular subtype.

The functional results and immune profiling that we found in autochthonous tumors and in human samples argue that *ARID1A* status intersects with immune surveillance in a manner that is not apparent in immune-naïve, *in vitro* based model systems. While our *in vivo* findings connect *ARID1A* to discrete steps of the cancer immunity cycle, the disparate immune phenotypes between tumor cells that had *Arid1a* deleted in *vitro* versus *in vivo* demonstrate that the immune-evasive program observed in *in-vivo-*R *Arid1a* KO tumors is not an obligate, immediate consequence of *Arid1a* deletion. Instead, our findings in preclinical models suggest that the immune consequences of tumor *Arid1a* loss are shaped by non-tumor cell-autonomous contexts. This hypothesis is supported by our findings in the TCGA dataset where *ARID1A*-mutant tumors had decreased IFN-γ pathway activity only in the GS subtype samples, and not in the CIN group.

Our current study builds upon previous work that demonstrated the importance of context when determining *ARID1A* functional effects in cancer.^8,21,35,36^ Specifically, in gastric adenocarcinoma, the molecular underpinning of the cancer is an essential context that must be considered as related to tumor *ARID1A* functionality. The TCGA identified four distinct molecular subtypes of gastric adenocarcinoma.^15^ *ARID1A* alterations are particularly enriched in EBV-positive and MSI subtypes, which are themselves associated with higher baseline immunogenicity and responsiveness to immune checkpoint blockade.^3^ In this study, we explored immune-related programs within the GS and CIN subtypes, which comprise the large majority of gastric adenocarcinomas, and found that GS *ARID1A* mutant tumors were immune-cold, consistent with our murine model, while CIN *ARID1A* mutant tumors were inflamed. Another recent study also considered how tumor-autonomous context determines *ARID1A* functionality by stratifying gastric cancers by Lauren histology into diffuse-type and intestinal-type, which are loosely associated with the GS and CIN subtypes, respectively.^15,37,38^ Diffuse-type gastric cancers have more frequent peritoneal metastases, are less sensitive to chemotherapy, and are associated with worse outcomes.^39,40^ The authors observed that *ARID1A* mutations in diffuse-type gastric cancer are associated with a poorer prognosis, but improved outcomes in intestinal-type tumors. In addition, the proteomes between the two groups were also markedly different.^37^ While the above study did not investigate the underlying mechanisms, it provides additional evidence that the effects of *ARID1A* mutations in gastric cancer are complex and the context in which these alterations occur may result in distinct or even opposing biological consequences.

Our finding that the immunogenic effect of tumor *ARID1A* loss is context-dependent provides an explanation for the contradictory conclusions in the literature regarding *ARID1A* functionality as related to cancer immunity.^3–8^ The disparate results in the literature may be due to differences across studies in various confounding factors that include, but are not limited to, tumor type, cooperating genomic alterations, timing of mutation acquisition, and the immune contexts (or lack thereof) in which the models form. A key implication of our data is that consideration of tumor *ARID1A* loss and immunogenicity must account for other variables. In support of this concept are recent studies indicating that *ARID1A*-associated immune phenotypes depend on cooperating oncogenic context, such as *BRAF^V^*^600^*^E^* in melanoma and *MYCN* amplification in neuroblastoma.^8,36^

An important direction for future research is to delineate the molecular mechanisms by which various contexts shape how tumor *ARID1A* interacts with host factors to determine the immunogenicity of developing or established malignancies. Prior work in multiple epithelial malignancies demonstrates that *ARID1A* loss promotes tumor progression and metastasis, which we also observed with *Arid1a* deletion regardless of whether its loss occurred *in vitro* or *in vivo*.^7,21,35^ However, tumor suppressor loss is not solely selected because it increases proliferative capacity. Tumor evolution occurs under multiple constraints, including immune surveillance, and accumulating evidence indicates that tumor suppressor alterations can also be selected for their ability to facilitate immune escape.^41^ In our study, the disparate immune phenotypes that we observed between the *in-vitro-*R and *in-vivo-*R systems may be considered using the immunoediting framework where tumor immunogenicity reflects the immunologic environment from which a cancer clone arises. Immune-mediated elimination and equilibrium impose selective constraints that ultimately favor outgrowth of clones capable of evading immune clearance.^13,14^ Within this framework, the developing *in-vivo-*R *Arid1a* KO tumors were “sculpted” by the host immune system to select for clones that suppressed GM-CSF and IFNGR1 production that facilitated immune evasion.

The heterogeneous and context-dependent effects of *ARID1A* mutations have direct implications for clinical translation as *ARID1A* mutation status is being evaluated as a stratification biomarker in immunotherapy trials. However, our data argue that *ARID1A* status alone is likely to be non-specific. Consistent with this thesis, we found that PD-1/PD-L1 blockade did not provide measurable benefit in the *Arid1a* KO mice. At the same time, our results indicate that *Arid1a*-deficient tumors are not intrinsically resistant to immune-mediated elimination. Rather, the immune-evasive state is associated with impaired initiation of the cancer immunity cycle, suggesting that approaches that provide strong innate activation and local immune priming may be effective. Consistent with this, a primed adaptive immune state reduced subsequent engraftment of *Arid1a* KO tumoroids relative to naïve hosts, indicating that shared tumor antigens can support clearance when a sufficiently robust immune response is induced. Furthermore, intratumoral administration of the diABZI STING agonist prevented engraftment and provided tumor control in *Arid1a* KO tumors, supporting a therapeutic strategy focused on restoring immune initiation in *ARID1A*-deficient, immune-excluded settings.

Several limitations of our study should be noted. First, the autochthonous tumors evolved with non-immune stromal compartments, such as cancer-associated fibroblasts, that shaped the immune microenvironment, while *in vitro* tumoroids develop in the absence of stromal input prior to transplantation. The relative contribution of the non-immune stromal components to the divergent phenotypes between the *in vivo* and *in vitro* systems therefore could not be fully resolved by the current study and should be a subject for future work. Second, since we performed immune profiling at early post-transplantation time points and at endpoint disease in autochthonous tumors, longitudinal analyses would refine the kinetics by which *Arid1a* status shapes immune recruitment versus exclusion. Finally, although we identify impaired GM-CSF output and attenuated IFN-γ responsiveness as major tumor-intrinsic mechanisms associated with *Arid1a* loss, these axes are unlikely to be exhaustive as *ARID1A* controls chromatin accessibility at a global level.

## CONCLUSION

In conclusion, our study identified immune selection pressure as a critical context that determined how tumor *ARID1A* interacted with the cancer immunity cycle. We showed that *ARID1A* loss was not sufficient to confer immune evasion but could promote immune escape after prolonged *in vivo* tumor development under immune pressure. Our findings argue that tumor *ARID1A* mutation status alone should not be interpreted as a uniform, pan-cancer determinant of immunogenicity or immune checkpoint blockade response without considering other contexts. More broadly, incorporation of immunoediting history into model design will be essential for defining the true immune consequences of tumor genomic alterations and for translating those insights into clinically actionable biomarker and therapeutic strategies.

## ACKNOWLEDGEMENTS

We acknowledge the assistance of the Tissue Management Shared Resource, the Quantitative Light Microscopy Core, and the Data Science Shared Resource, all of which are shared resources at the UT Southwestern Harold C. Simmons Comprehensive Cancer Center and supported by the National Cancer Institute (NCI; P30CA142543). We also acknowledge the UT Southwestern Whole Brain Microscopy Facility (RRID: SCR_017949), for use of whole slide scanning resources.

R.T.H. and J.D.K. were supported by UT Southwestern Training Resident Doctors as Innovators in Science awards from the Burroughs Wellcome Fund (1018897). M.F.P. was supported by the NCI (3R37CA265967-01A1S1) and the American College of Surgeons Resident Research Scholarship. S.L.N. was supported by the UT Southwestern M.D. Scientist Training Program and Society for Surgery of the Alimentary Tract Residents and Fellows Research Award. T.H.H. was supported by the NCI (R01CA276690) and from the Department of Defense (CA190578), Schmidt Sciences through the Mayo Clinic Foundation, and the Torrey Coast Foundation. S.T.G.H. was supported by the NCI (R37CA265967, R01CA276690). H.Z. was supported by the Mark Foundation for Cancer Research (21-003-ELA), the Simmons Comprehensive Cancer Center (P30CA142543), the Cancer Prevention and Research Institute of Texas (RP220614), and the NIH (DP1DK139976, R01CA251928). I.S.C. was supported by the Susan G. Komen Career Catalyst Award (1010879) and the NCI (K08CA270188). S.C.W. is a UT Southwestern Disease Oriented Clinical Scholar and was supported by NCI (R37CA265967, R01CA276690).

## DECLARATION OF INTERESTS

T.H.H. is a co-founder of Kure.ai therapeutics and has received consulting fees from IQVIA; these affiliations and financial compensations are unrelated to the current paper. H.Z. is a co-founder of Quotient Therapeutics and Jumble Therapeutics and serves as an advisor for Newlimit; these affiliations are unrelated to the current paper.

## MATERIALS AND METHODS

### Animals

All animal experiments were approved by the Institutional Animal Care and Use Committee (IACUC) at the University of Texas (UT) Southwestern Medical Center and conducted in accordance with institutional guidelines. Mice were maintained under specific pathogen-free conditions on a 12-hour light/dark cycle at 20–26 °C and 30–70% relative humidity, with food and water provided *ad libitum*. All mice were maintained on a C57BL/6 background (backcrossed ≥6 generations). Mice carrying floxed alleles of *Trp53* (#008462), *Cdh1* (005319), and the *Rosa26*^LSL-eYFP^ reporter allele (006148) were obtained from The Jackson Laboratory. Mice carrying floxed *Arid1a* alleles were kindly provided by Dr. Hao Zhu (UT Southwestern Medical Center). Mice carrying the *Atp4b-Cre* allele were kindly provided by Dr. Sam Yoon (Columbia University). Experimental cohorts were generated by intercrossing *Trp53*^flox/flox^; *Cdh1*^flox/flox^; *Rosa26*^LSL-eYFP/LSL-eYFP^; *Arid1a*^+/flox^ parents to ensure all experimental animals were littermate controls. Resulting offspring carried *Atp4b-Cre*; *Trp53*^flox/flox^; *Cdh1*^flox/flox^; *Rosa26*^LSL-eYFP/LSL-eYFP^ and differed exclusively in *Arid1a* zygosity: *Arid1a*^+/+^ (wild-type), *Arid1a*^+/flox^ (heterozygous), or *Arid1a*^flox/flox^ (knockout). All analyses were performed in an age- and sex-matched manner unless otherwise indicated in the figure legends.

Genotyping was performed using the KAPA2G Fast HS Genotyping Mix (7961316001, Roche Diagnostics) in a total reaction volume of 11 μL containing 5 μl KAPA2G Fast HS Genotyping Mix, 4 μL water, 0.5 μL forward primer (5 μM), 0.5 μL reverse primer (5 μM), and 1 μL template DNA. PCR cycling conditions were identical for all alleles: initial denaturation at 95°C for 3 min; 30 cycles of 95°C for 15 s, 58°C for 20 s, and 72°C for 45 s; followed by a final extension at 72°C for 5 min. PCR products were resolved on a 2% agarose gel in 1× TAE buffer containing ethidium bromide and run at 200V for 40–45 min. All primers and expected band sizes are listed in the resource table.

### Tumoroid culture media formulation

The *<u>basal medium</u>* was prepared using Advanced Dulbecco’s modified Eagle’s medium/F12 (12634-028, Invitrogen) and supplemented with HEPES (10 mM; 15630-080, Invitrogen), GlutaMAX (1X; 35050-061, Invitrogen), penicillin/streptomycin (100 U/mL / 100 μg/mL; 15140-148, Invitrogen), N2 supplement (1x; 17502-048, Invitrogen), B27 supplement (1X; 17504-044, Invitrogen), N-acetylcysteine (1 mM; A9165, Sigma-Aldrich), and bovine serum albumin, Fraction V (1%; BP1600, Fisher Scientific)

The *complete medium* was prepared by supplementing the basal medium with noggin: 100 ng/mL (HY-P70558, MedChemExpress), epidermal growth factor: 50 ng/mL (HY-P700051AF, MedChemExpress), fibroblast growth factor 10: 100 ng/mL (HY-P7048, MedChemExpress), R-spondin 1: 100 ng/mL (HY-P7114, MedChemExpress), Wnt3a: 100 ng/mL (HY-P70453B, MedChemExpress), gastrin I: 10 nM (HY-P1097, MedChemExpress), and Y-27632 ROCK inhibitor: 10 µM (ATCC, ACS3030).

### Establishing cell lines from mouse tumors

Tumors were harvested from mice stomachs and washed three times in ice-cold phosphate-buffered saline (PBS) containing 1× penicillin-streptomycin. The tumor was minced to pieces smaller than 1-2 mm³ and incubated with enzymatic cocktail comprised of 2 mg/mL collagenase type IV and 0.1 mg/mL DNase I in PBS. The mixture was incubated at 37 °C for 1 hour with inversion every 10 minutes. The reaction was stopped by adding an equal volume of DMEM/F12 supplemented with 10% fetal bovine serum (FBS) and 1% penicillin-streptomycin, and the mixture was passed through a 70-µm cell strainer. The cell suspension was centrifuged at 300 × g for 5 minutes at room temperature, the supernatant was aspirated, and the pellet was resuspended in 5 mL of DMEM/F12 supplemented with 10% FBS and 1% penicillin-streptomycin. Cells were seeded into a 10-cm dish and cultured at 37 °C in 5% CO₂. Cells were routinely passaged every three days using TrypLE Express (12604013, Thermo Fisher Scientific) dissociation and re-seeded at the desired density for maintenance.

### Establishing tumoroids from mouse tumors

Tumors were harvested from mice stomachs and washed three times in ice-cold PBS with 1% penicillin-streptomycin. The tissue was minced into 1–2 mm³ pieces and incubated in a digestion buffer containing collagenase type IV and DNase I dissolved in Advanced DMEM/F12. Digestion was performed at 37°C for 1 hour with gentle shaking. The suspension was filtered through a 70-µm cell strainer, centrifuged at 300 x g for 5 minutes at 4°C, and washed once with cold basal medium. Tumoroids were resuspended in 50 µL domes of Growth Factor Reduced Matrigel (356230, Corning) in pre-warmed 48-well plates. After polymerization, 200 µL of complete medium was added. Medium was changed every 2–3 days. Tumoroids were passaged every 5–7 days at a ratio of 1:5. For passaging, the culture medium was removed, and ice-cold PBS was added to dissolve the Matrigel. The tumoroid suspension was transferred to a 15 mL conical tube. Mechanical dissociation was performed by pipetting the suspension using a fire-polished glass Pasteur pipette. Care was taken to avoid excessive pipetting to prevent dissociation into single cells. The fragments were centrifuged at 500 x g for 5 minutes at 4°C. The supernatant was discarded, and the pellet was resuspended in fresh Matrigel for re-plating and cultured at 37°C in 5% CO₂.

To ensure the genetic purity of the tumor cells, tumoroids were subjected to selection by nutlin-3a (SML0580, MilliporeSigma), which was added to the culture medium at 10 µM for a period of 14 days. Following this selection phase, cultures were examined using fluorescence microscopy. Only lines confirmed to be 100% YFP-positive were selected and maintained for subsequent experiments.

### Flank injections

Tumor cells were harvested via 0.25% trypsin-EDTA and were then counted and resuspended in DMEM supplemented with 10% FBS and 1% penicillin/streptomycin. 1,000,000 cells (total volume 100 µL) were injected. Tumoroids were expanded, harvested, and washed three times with cold PBS. Cells were dissociated to single cells using TrypLE Express, counted, and resuspended in basal medium mixed 1:1 with Matrigel. 300,000 cells (total volume 200 µL) were injected. Tumors were measured with calipers, and tumor volumes were calculated using the formula: (π/6) x length x width^2^. Mice were monitored until tumors reached institutional endpoint criteria or pre-determined endpoint.

### Tail vein injections

Tumor cells were harvested via 0.25% trypsin-EDTA and filtered through a 40 μm strainer and resuspended in cell media. 1,000,000 viable cells in 100 µL of complete media were injected via the lateral tail vein using a 28-gauge needle. Tumoroid lines were harvested from Matrigel and dissociated with 0.25% trypsin-EDTA. They were then filtered through a 40 μm strainer and resuspended in complete media. 300,000 viable cells in 100 μL of complete media were injected via the lateral tail vein using a 28-gauge needle. Mice were monitored for health and euthanized 2–8 weeks later for lung collection and histologic analysis.

### Immunohistochemistry

Paraffin-embedded tissue sections were deparaffinized in xylene, rehydrated through a graded ethanol series, and subjected to antigen retrieval using 10% Citra buffer (HK0809K, Biogenex Laboratories) microwaved for 20 minutes in a water bath. After cooling, slides were quenched in 3% H₂O₂ in methanol, blocked with 1.5% serum in PBS-Tween for 1 hour, and incubated overnight at 4°C with primary antibody. The next day, slides were washed and incubated with biotinylated secondary antibody, followed by incubation with VECTASTAIN® Elite ABC-HRP Kit (PK6101 and 6104, Vector Laboratories) for 30 minutes. DAB (SK-4100, Vector Laboratories) was used as the chromogen, and slides were counterstained with hematoxylin, rinsed, and mounted with permanent mounting media.

### Immunofluorescence

Paraffin-embedded tissue sections were deparaffinized in xylene, rehydrated through a graded ethanol series, and subjected to antigen retrieval using 10% Citra buffer microwaved for 20 minutes in a water bath. After cooling, slides were quenched in 3% H₂O₂ in methanol, blocked with 5% bovine serum albumin in PBS-Tween for 1 hour, and incubated overnight at 4°C with primary antibody. The following day, slides were washed and incubated with a fluorophore-conjugated secondary antibody for 1 hour at room temperature in the dark. After washing, slides were mounted using a DAPI-containing antifade mounting medium (P36935, Invitrogen) and stored at 4°C protected from light until imaging.

### Overexpressing *ARID1A* in cell lines

*Arid1a* knockout cell lines were transfected with the PT3-EF1a-hArid1a-T2A-Puromycin plasmid using Lipofectamine 3000 (L3000008, Thermo Fisher Scientific). Formulated lipid-DNA complexes were added to cells at 70% confluency. Forty-eight hours post-transfection, stable transformants were selected by adding 2 µg/mL of puromycin to the culture medium. Following selection, single-cell clones were isolated via limiting dilution and expanded.

### *In vitro* induction of gene mutations

To induce Cre-mediated gene recombination, organoids were dissociated into small clusters using TrypLE Express. The cell suspension was incubated with Adenovirus-Cre (VVC-U of Iowa-5 Ad5CMVCre, Viral Vector Core, University of Iowa, USA) supernatant. The cells were reseeded in Matrigel and cultured in complete medium. One week post-transduction, cultures were checked for YFP expression by fluorescence microscopy.

### Exogenous interferon-treatment of tumoroids

Tumoroids were seeded and cultured for 24 hours under standard conditions. Culture medium was removed and replaced with fresh complete medium supplemented with recombinant mouse IFN-γ (575304, Biolegend) at 20 ng/mL. The tumoroids were incubated at 37°C for 24 hours and then analyzed.

### Flow cytometry of tumoroids

Tumoroid lines were harvested from Matrigel and dissociated with 0.25% trypsin-EDTA. They were then filtered through a 40 μm strainer and centrifuged at 300x g for 7 minutes. Pellets were resuspended in blocking buffer (1 μL rat anti-mouse CD16/32 (553142, BD Pharmingen) per 70 μL staining buffer (00-4222-26, Invitrogen)) and incubated at 4°C for 15 minutes. Primary antibody and fixable viability dye (L34992, Thermo Fisher Scientific) diluted in staining buffer was added to each tube, and cells were incubated for 1 hour at 4°C in the dark. After washing, cells were fixed in 150 μL of fixation buffer (00-8222-49, Invitrogen) and stored at 4°C protected from light until analysis (within 72 hours).

### Immune cell isolation and co-culture establishment

Spleens were harvested from mice and dissociated using the Spleen Dissociation Kit (130-095-926, Miltenyi Biotec). Dendritic cells and CD8^+^ T cells were isolated using the Pan Dendritic Cell Isolation Kit (130-100-875, Miltenyi Biotec) and CD8a^+^ T Cell Isolation Kit (130-104-075, Miltenyi Biotec), respectively. For tumoroid preparation, a split-sample quantification strategy was used to preserve 3D structure: one aliquot was enzymatically digested to single cells for counting, while the matching aliquot (containing tumoroid fragments) was used for seeding. Tumoroid fragments were mixed with immune cells at a ratio of 1:200, resuspended in RPMI 1640 + 10% FBS, and mixed with Matrigel (final concentration 30%). Co-cultures were maintained in RPMI 1640 + 10% FBS.

### *In vivo* immune cell depletion

Mice received intraperitoneal injections of depleting antibodies (200 µg/injection) starting 3 days prior to tumor challenge and continuing every 3 days until Day 18. The following antibodies were used (BioXCell): anti-CD8a (Clone 53-6.7), anti-CD4 (Clone GK1.5). Controls: rat IgG2a (Clone 2A3) and rat IgG2b (Clone LTF-2). Depletion was confirmed by flow cytometry of peripheral blood.

### Microscopic imaging and quantification of dendritic cell recruitment

Dendritic cells were labeled with CellTracker™ Deep Red (C34565, Thermo Fisher Scientific) in RPMI 1640 for 1 hour at 37°C prior to co-culture. Tumoroids were identified by their intrinsic YFP expression, and dendritic cells were visualized via the CellTracker™ Deep Red fluorescence. Dendritic cell and tumoroid co-cultures were imaged using a fluorescence microscope Leica DMi8. Images were acquired from random fields of view for each condition. The degree of interaction was determined by counting the number of dendritic cells co-localized with individual tumoroids.

### diABZI administration

diABZI (S8796, Selleck Chemicals) was prepared fresh each day in 5% DMSO and 95% corn oil on at a dose of 1.5 mg/kg at a final volume of 100 µL. For treatment of established tumors, diABZI was injected into the flank tumors at days 21, 28, and 35. Tumors were harvested for flow cytometry 72 hours following the last injection. For diABZI treatment of newly-injected tumoroids, 100 µL of diABZI was injected into the marked flanks 1, 7, and 14 days after tumoroids injection.

#### Flow cytometry of flank tumors

Subcutaneous tumors were harvested, washed in cold PBS, and then dissociated into single-cell suspensions using the Tumor Dissociation Kit (130-096-730, Miltenyi Biotec), passed through a 40 µm cell strainer, washed with cold staining buffer, and incubated with anti-mouse CD16/CD32 (553142, BD Pharmingen) for 10 minutes at 4°C. The specific fluorophore-conjugated antibodies were added to the cell suspension. The samples were incubated for 1 hour at 4°C in the dark. Following incubation, cells were washed twice with cold staining buffer. For overnight storage, the final cell pellets were resuspended in 200 µL of a 1:1 mixture of Flow Cytometry Staining Buffer (00-4222-26, Thermo Fisher Scientific) and IC Fixation Buffer (00-8222-49, Thermo Fisher Scientific). The samples were stored at 4°C in the dark overnight, and data acquisition was performed the following day on an Attune NxT Flow Cytometer or the Aurora Spectral flow cytometer, depending on the assay. The acquired data were analyzed using FlowJo (version 11).

### Cytokine analysis

Conditioned media were harvested from tumoroid cultures following a 48-hour incubation period. All harvesting and processing steps were performed on ice. The entire well contents, comprising both the culture medium and the Matrigel dome, were collected and mechanically disrupted using wide-bore pipette tips. The suspension was centrifuged at 4°C and supernatant was filtered through a 40 µm cell strainer and stored at -70°C. Cytokine levels were quantified using the Mouse Cytokine 32-Plex Discovery Assay (Eve Technologies, Calgary, AB, Canada). To normalize cytokine secretion data, the cell pellet from the centrifugation step was dissociated into single cells using TrypLE Express and counted. Final cytokine concentrations were calculated by normalizing the results to the total cell number per well.

### ATAC-sequencing preparation

Tumoroids were harvested and washed with ice-cold PBS. 100,000 cells were processed using the ATAC-Seq Kit (53150, Active Motif). Nuclei were permeabilized and confirmed by Trypan Blue staining and tagmented. Libraries were sequenced on an Illumina NextSeq 2000.

### Tumor bulk RNA sequencing preparation

Snap-frozen tumor tissue collected from mice was homogenized in TRIzol™ using a bead mill. RNA was purified using the RNeasy Mini Kit (74104, Qiagen). Libraries were constructed using the NEBNext® Ultra™ II Directional RNA Library Prep Kit (E7760L, New England Biolabs) and sequenced on an Illumina MiniSeq. Tumoroids were harvested from culture and washed three times with ice-cold PBS and the tumoroids were resuspended in TRIzol™ Reagent (15596026, Thermo Fisher Scientific) and homogenized by vortexing. RNA was purified using the RNeasy Mini Kit and sequenced on an Illumina NovaSeq.

### Tumor single-cell RNA sequencing preparation

Stomach tumors were harvested from mice and washed in ice-cold PBS. The tissue was dissociated into single-cell suspensions using the Tumor Dissociation Kit (130-096-730, Miltenyi Biotec) and passed through a 70-µm cell strainer and debris was removed using the Debris Removal Solution (130-109-398, Miltenyi Biotec). Single-cell suspensions were chemically fixed using the Fixation Kit (ECFC3300, Parse Biosciences). Fixed cells were stored at -70°C until library preparation. For library construction, samples were thawed, and barcoding was performed using the Evercode™ Whole Transcriptome Kit (ECWT3300, Parse Biosciences).

Cells were pooled, redistributed, and subjected to subsequent rounds of ligation to append unique combinatorial barcodes. Following barcoding, cells were lysed, and the library was amplified and purified. The final library concentration and size distribution were assessed using Agilent TapeStation. The resulting scRNA-seq libraries were sequenced on an Illumina NextSeq 2000 system using paired-end sequencing protocols compatible with the Parse Biosciences pipeline. A 5% PhiX control library spike-in was added to the sequencing run to ensure library diversity.

### Quantitative reverse-transcription PCR

Tumoroids were harvested on ice. The dome and medium were collected, centrifuged, and washed twice with cold PBS before lysis in TRIzol™. RNA was purified using the RNeasy Mini Kit (Qiagen). cDNA was synthesized using the LunaScript® RT SuperMix Kit (E3010, New England Biolabs). A total of 2 µg of RNA was used per 40 µL reaction. The reverse transcription reaction was carried out in a thermocycler with the following conditions: primer annealing: 25°C for 2 minutes, cDNA synthesis: 55°C for 10 minutes, heat inactivation: 95°C for 1 minute. The resulting cDNA was used as a template for qPCR analysis. qPCR was performed using SYBR Green Master Mix (1708880EDU, Bio-Rad) and run on a QuantStudio 3 system (Thermo Fisher Scientific). Relative gene expression was calculated using the ΔΔCt method.

### Statistical analyses

Data are presented as mean ± SEM unless otherwise indicated, and the statistical test for each panel is specified in the corresponding figure legend. GraphPad Prism (version 11) was utilized.

### ATAC-seq data analysis

Raw read quality was assessed with FastQC (v0.11.9)^1^. Adapters and low-quality bases were removed in paired-end mode with Trim Galore (v0.6.7)^2^. Reads were aligned to the mouse reference genome (UCSC mm10) with BWA-MEM (v0.7.17)^3^, and secondary alignments were removed with SAMtools (v1.16.1)^4^. PCR duplicates were marked with Picard MarkDuplicates (v3.0.0)^5^. Aligned reads were then filtered with BamTools (v2.5.2)^6^ and SAMtools (v1.16.1)^4^ to remove unmapped or multi-mapped reads (MAPQ < 1), improperly paired reads, soft-clipped reads, reads with excessive mismatches (>4), read pairs with abnormally large insert sizes (>2 kb), and reads overlapping regions in the ENCODE mm10 blacklist v2^7^ or mitochondrial DNA (chrM). To address the 9-bp duplication introduced by Tn5 transposition, the filtered alignments were shifted with deepTools (v3.5.6; alignmentSieve --ATACshift)^8^. Open-chromatin narrow peaks were then called per sample on the shifted alignments with MACS2 (v2.2.9.1; --format BAMPE --nomodel --shift -100 --extsize 200 --keep-dup all)^9^. Per-sample ATAC-seq quality metrics, including TSS enrichment and fragment-size distribution, were assessed with ataqv (v1.3.1)^10^. Differential accessibility analysis was performed with DiffBind (v3.12.0)^11,12^ in R (v4.3.3). Differential peaks were then annotated to the nearest gene with ChIPpeakAnno (v3.36.0)^13,14^. Locus-level chromatin-accessibility tracks at candidate genes were visualized with Gviz (v1.46.1)^15^.

### Tumoroid RNA sequencing data analysis

Adapters and low-quality bases were trimmed from paired-end RNA-seq reads with Trim Galore (v0.6.4)^2^. Trimmed reads were aligned to the mouse reference genome (UCSC mm10) with STAR (v2.4.2a; --alignSJoverhangMin 8 --alignSJDBoverhangMin 1 --alignIntronMin 20 -- alignIntronMax 1000000 --alignMatesGapMax 1000000)^16^. PCR duplicates were marked and removed with Picard MarkDuplicates (v2.10.3)^5^ and SAMtools (v1.8)^4^. Gene-level read counts were quantified in un-stranded mode with Subread featureCounts (v1.6.1)^17^. Differential expression analysis (knockout vs. wild-type) was performed with DESeq2 (v1.42.0)^18^ in R (v4.3.3). Gene-set enrichment analysis (GSEA) was performed with the fgsea package^19^ (v1.28.0) against the MSigDB mouse Hallmark collection.^20^

### Mouse tumor bulk RNA sequencing data analysis

Gastric cancer RNA-seq libraries were quantified against the mouse reference transcriptome (GRCm39) with Salmon^21^ and summarized to gene level using tximport.^22^ Gene-level counts were analyzed with DESeq2^18^ (design ∼ batch + genotype, across three sequencing batches); sample sex was excluded from the model owing to confounding with genotype. Genes with CPM > 1 in at least as many samples as the smallest genotype group were retained. Differential expression between homozygous *Arid1a* knockout (Homo) and wild-type (WT) tumors was assessed using the Wald test with WT as reference and Benjamini–Hochberg correction, with adjusted p < 0.05 considered significant. Variance-stabilizing-transformed counts were used for principal-component and sample-distance quality control, and Ensembl identifiers were annotated with biomaRt.^23^

### Gene set enrichment analysis (mouse)

Genes were ranked by the DESeq2^18^ Wald statistic (duplicate symbols collapsed to the largest absolute value) and tested against the MSigDB mouse Hallmark collection^20^ using fgsea^19^ (gene sets of 15–500 genes, 1,000 permutations), with normalized enrichment scores and Benjamini– Hochberg–adjusted p-values reported.

### Gene set enrichment analysis of TCGA-STAD samples

RNA-seq expression and somatic mutation data from TCGA STAD PanCancer Atlas cohort (stad_tcga_pan_can_atlas_2018) were obtained via the cBioPortal^24,25^ REST API, and patients with documented TCGA subtypes of genomically stable (GS) and chromosomal instability (CIN) subtypes were extracted. *ARID1A* loss-of-function (LOF) was defined from somatic mutation calls (Entrez 8289) as a nonsense, frameshift, or splice-site/splice-region mutation, with all remaining tumors in each subtype designated wild-type (WT), yielding 6 LOF/44 WT (GS) and 16 LOF/207 WT (CIN). Within each subtype, genes were ranked between LOF and WT, computed on the log2 RSEM values provided by cBioPortal. Pre-ranked GSEA against the MSigDB Hallmark collection^20^ (50 gene sets) was performed with fgsea^19^ (gene sets of 15–500 genes, 1,000 permutations, fixed random seed), with FDR < 0.05 considered significant.

### Cross-species molecular subtype classification

Mouse bulk RNA-seq profiles were projected onto the four TCGA molecular subtypes of gastric adenocarcinoma (EBV, MSI, CIN, GS) using 13,072 one-to-one mouse–human orthologs (Ensembl BioMart^23^). A Prediction Analysis of Microarrays (PAM) nearest shrunken centroid classifier^26^ was trained on 274 TCGA STAD bulk RNA primary tumors with documented TCGA subtypes, using the top 100 markers per subtype selected by limma^27^ one-versus-rest differential expression (FDR < 0.05, log2 fold change > 0.25). The classifier was applied to 29 samples from *in-vivo-*R mice stomach tissue (WT n = 11, Het n = 7, KO n = 11) and subsequently predicted each subtype. The genotype–subtype association was evaluated by Fisher’s exact test and a 1,000-permutation genotype-label test.

### Single cell RNA sequencing

#### Library Prep and Preprocessing

Single cell RNA-seq libraries were prepared using Parse Biosciences split-pool combinatorial barcoding chemistry^28^ from gastric tissue of *in-vivo-*R *Arid1a* WT (N = 4) and *in-vivo-*R *Arid1a* KO (N = 4) mice. Libraries were sequenced on the Singular Genomics G4 platform. Raw reads were processed through the Parse Biosciences Trailmaker pipeline aligned to the mouse genome GRCm38 to generate per-cell gene expression matrices. Gene identifiers were retained as Ensembl IDs (ENSMUSG format) throughout all analyses and converted to gene symbols via org.Mm.eg.db^29^ (Bioconductor 3.18) at the reporting stage. The Trailmaker-exported Seurat^30^ object contained 36,239 cells across the 8 samples.

Quality control filtered cells with nFeature_RNA >= 200, percent.mt <= 20, doublet scores <= 0.4. Sample_F6 (*in-vivo-*R *Arid1a* KO, female) was excluded due to sample-level technical failure: median nFeature_RNA of 257 (vs. cohort range 325-548 for other samples). The final primary atlas comprised 22,836 cells across 7 samples (4 *in-vivo-*R *Arid1a* WT: M1, M6, sample_F3, sample_F4; 3 *in-vivo-*R *Arid1a* KO: M3, M5, F2_6) after quality control metrics.

To augment cell-type resolution secondary to our modest sequencing depth (median 483 genes/cell, median 1,315 UMIs/cell) — we integrated 18,930 cells from an independently generated gastric single cell RNA-seq reference dataset from the same mouse genetic background (*Atp4b-Cre; Cdh1^flox/flox^; Trp53^flox/flox^*). These reference samples were processed on the same Parse Biosciences + Singular G4 platform and exhibited substantially higher per-cell gene detection (median 1,069-1,770 genes/cell across samples), providing a high-resolution cell-type identity scaffold.

Quality control for the reference dataset was matched to the experimental cohort (nFeature_RNA >= 200, percent.mt <= 20, doublet score <= 0.4). These samples were not included in the final analysis and served as anchors to help with cell-type resolution.

#### Normalization and integration

Experimental and Reference datasets were merged, which provided 41,766 cells across ten samples. Normalization was performed using SCTransform^31,32^ (sctransform v0.4.2) with the glmGamPoi^33^ method for accelerated parameter estimation. Batch correction was performed using Harmony^34^ (v1.2.4) on the sample variable (7 AridSingular + 3 reference samples), with theta=2 and the top 30 principal components as input after evaluating elbow plots. Integration quality was assessed using silhouette analysis on a 5,000-cell subsample.

UMAP^35^ dimensional reduction was computed from the Harmony-corrected embedding (30 dimensions). Shared nearest-neighbor graph construction and Louvain^36^ clustering were performed at four resolutions (0.3, 0.5, 0.7, 1.0) to assess cluster stability. Resolution 0.5 (20 clusters) was selected as the primary working resolution based on cluster coherence and biological interpretability.

#### Cell type annotations

Annotation was performed initially of broad compartments (Epithelial, Immune, Stromal). For each of the 20 clusters at resolution 0.5, differentially expressed genes were identified using Seurat’s^30^ FindMarkers (v5.0.3) (Wilcoxon rank-sum test, SCT assay, min.pct = 0.1, logfc.threshold = 0.25) against all other clusters. Gene symbols were mapped from Ensembl IDs via org.Mm.eg.db.^29^ Module scores (Seurat AddModuleScore) were computed for 27 canonical cell-type marker panels spanning gastric epithelial lineage, immune populations (T cell: *Cd3e, Cd3d*; B cell: *Cd19, Cd79a*; myeloid: *Itgam, Cd68, Lyz2*; plasma: *Jchain, Mzb1*), and stromal cells (fibroblast: *Col1a1, Dcn, Pdgfra*; endothelial: *Pecam1, Cdh5*). Mean module scores per cluster were used to generate proposed annotations, with confidence assessed by the margin between the top- and second-scoring panels. Twelve clusters were assigned to the Epithelial compartment, six to Immune, and two to Stromal. Due to modest resolution, the Epithelial clusters were collapsed to one single compartment.

All single cell analyses were performed in R (v4.3.1) with Seurat (v5.0.3), sctransform (v0.4.2), Harmony (v1.2.4), glmGamPoi (v1.14.3), and org.Mm.eg.db (v3.16.0, Bioconductor 3.18) on Red Hat Enterprise Linux 9.4 (BioHPC cluster, UT Southwestern). Complete analysis scripts for each stage will be deposited to an online repository. Key parameter choices (QC thresholds, Harmony theta, clustering resolutions) were evaluated through systematic sensitivity analyses.

---

**Supplemental Figure 1.**
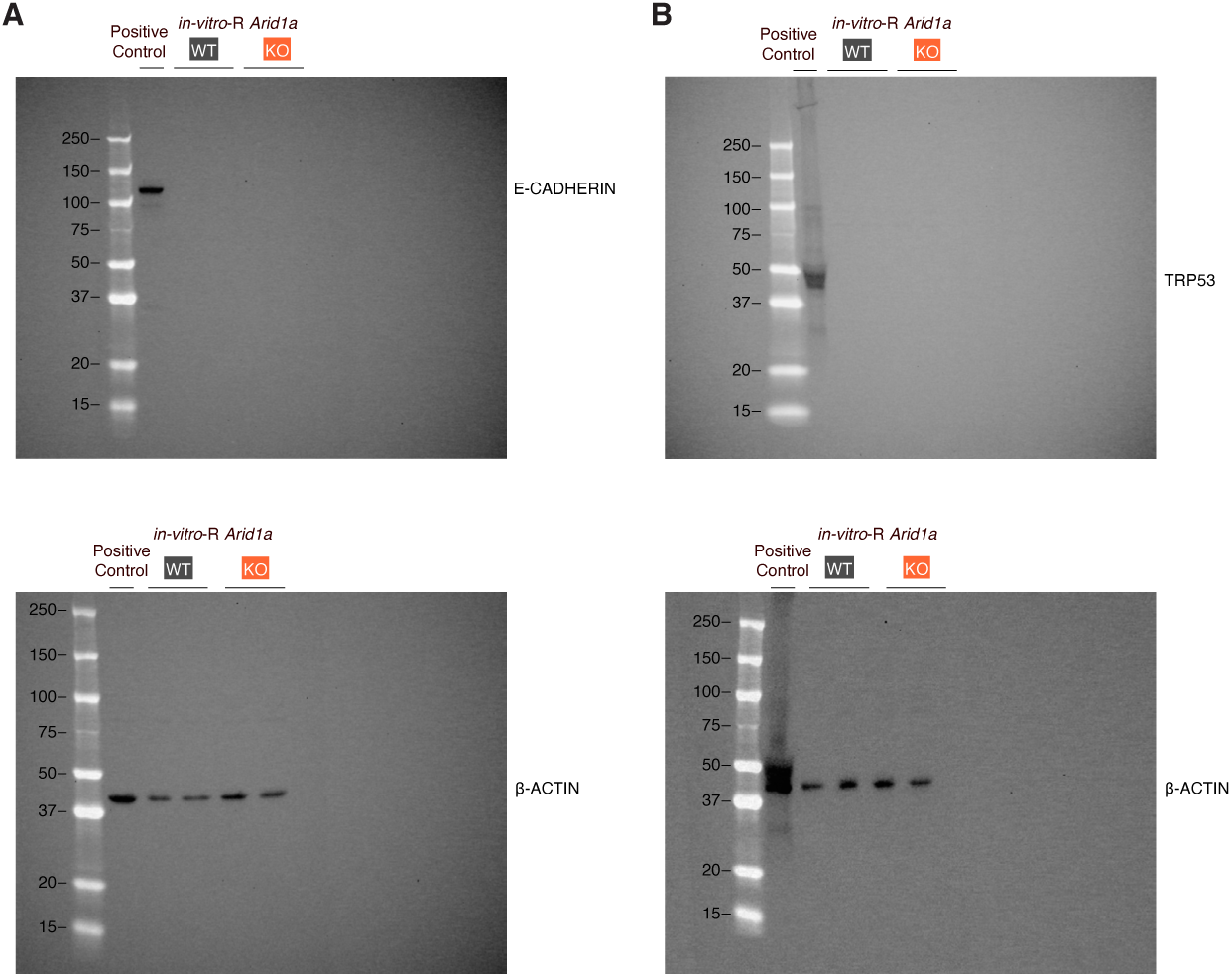
Related to Figure 1. Full Western blots of *in-vitro*-R *Arid1a* WT and KO tumoroids after Cre recombination, probed for **(A)** E-CADHERIN and β-ACTIN and **(B)** TRP53 and β-ACTIN.

**Supplemental Figure 2.**
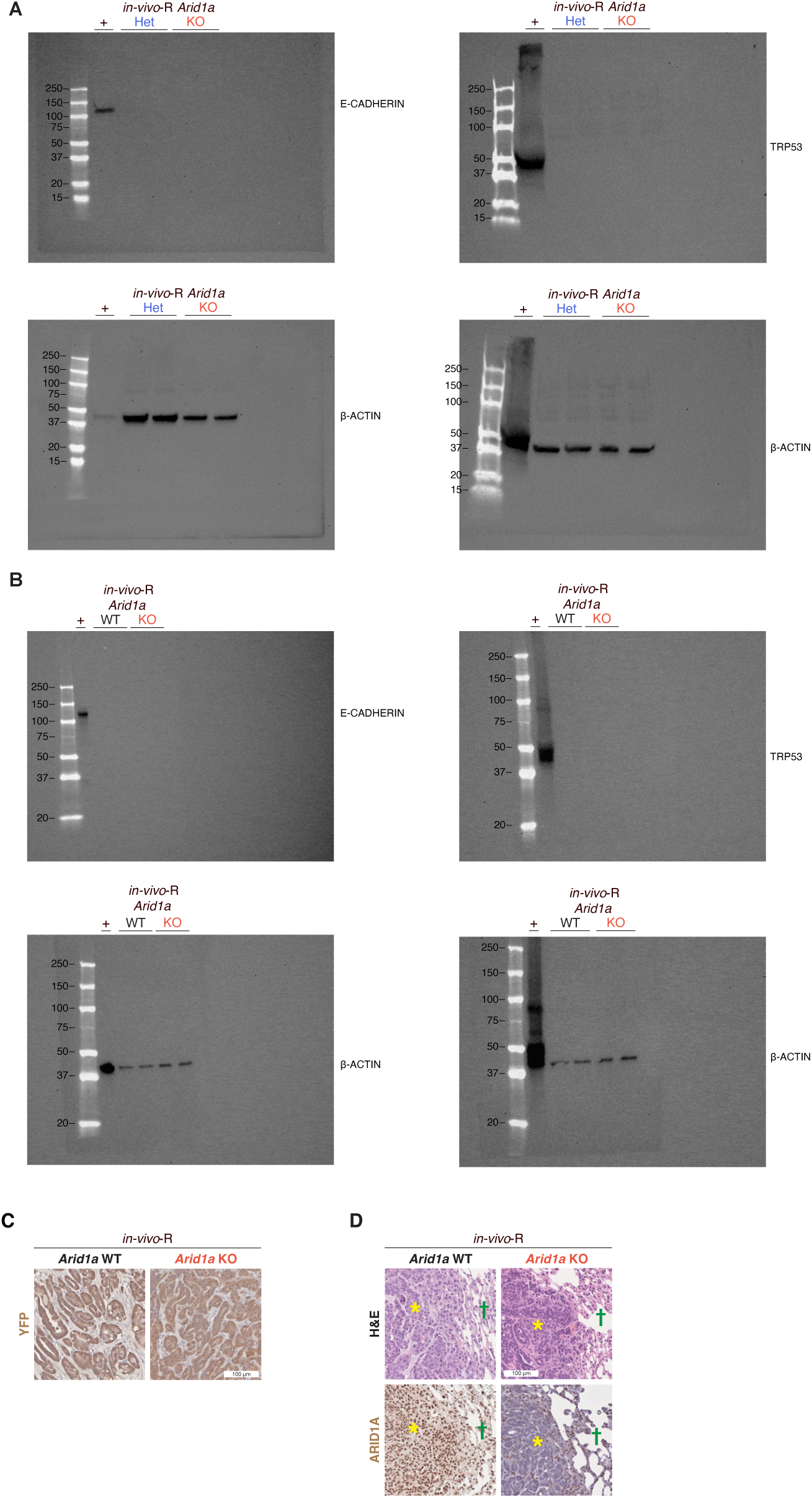
Related to Figure 2. **(A)** Full Western blots of *in-vivo-*R *Arid1a* cell lines probed for E-CADHERIN, TRP53, and β-ACTIN. **(B)** Full Western blots of *in-vivo-*R *Arid1a* tumoroids probed for E-CADHERIN, TRP53, and β-ACTIN. **(C)** YFP immunohistochemistry of tumors from NSG mice injected with *in-vivo-*R *Arid1a* WT and KO tumoroids. **(D)** Representative histologic images of lung metastases following tail vein injection of *in-vivo-*R *Arid1a* WT and KO tumoroids into NSG mice. Yellow stars indicate carcinoma; green crosses indicate adjacent normal lung.

**Supplemental Figure 3.**
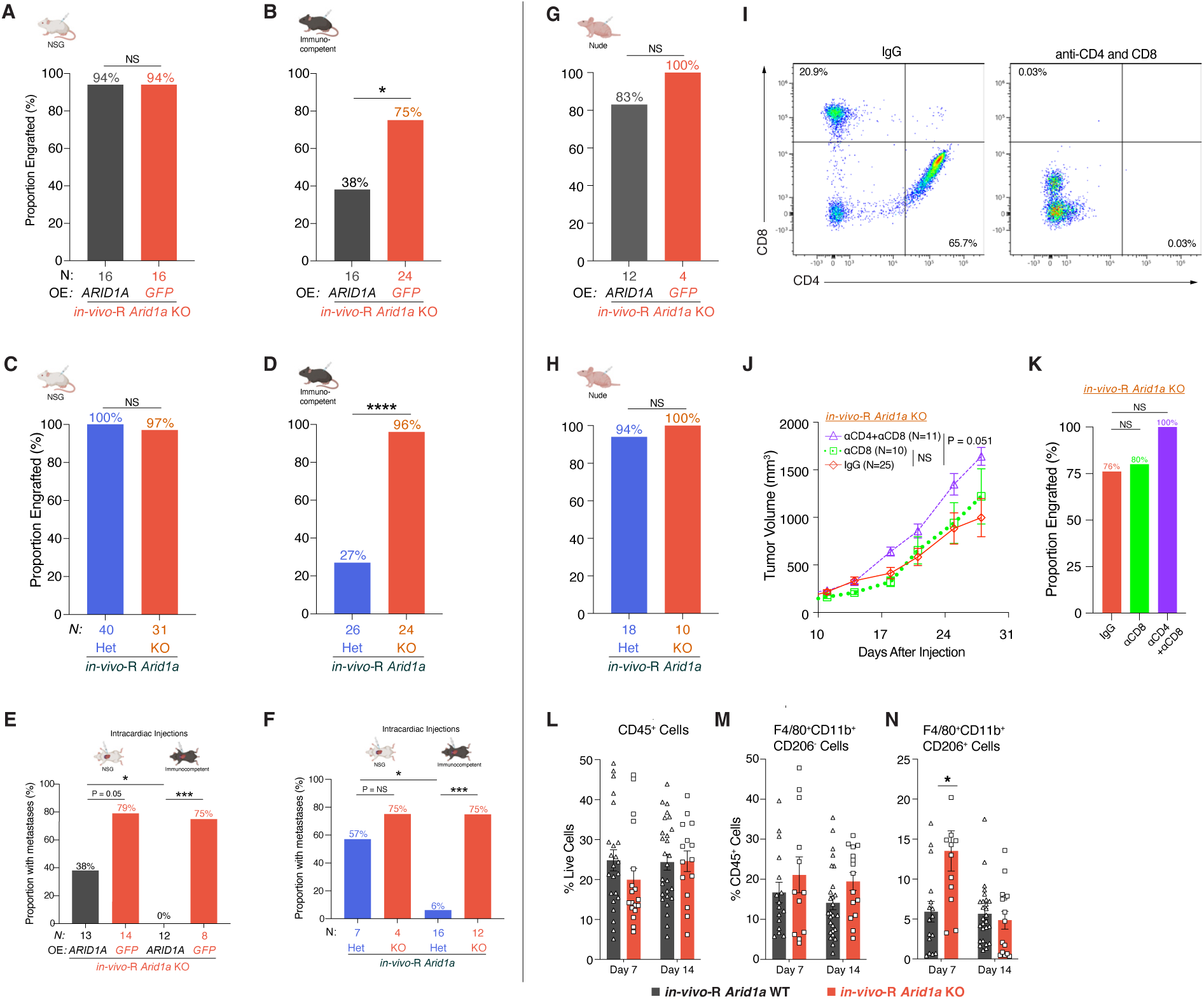
Related to Figure 3. **(A)** Proportion of NSG mice that developed tumors following flank injection of *in-vivo-*R *Arid1a* KO cells overexpressing *ARID1A* or *GFP*. **(B)** Proportion of syngeneic immunocompetent mice that developed tumors following flank injection of *in-vivo-*R *Arid1a* KO cells overexpressing *ARID1A* or *GFP*. **(C)** Proportion of NSG mice that developed tumors following flank injection of *in-vivo-*R *Arid1a* Het or KO cell lines. **(D)** Proportion of syngeneic immunocompetent mice that developed tumors following flank injection of *in-vivo-*R *Arid1a* Het or KO cell lines. **(E)** Proportion of NSG and syngeneic immunocompetent mice that developed metastases following intracardiac injection of *in-vivo-*R *Arid1a* KO cells overexpressing *GFP* or *ARID1A*. **(F)** Proportion of NSG and syngeneic immunocompetent mice that developed metastases following intracardiac injection of *in-vivo-*R *Arid1a* Het or KO cell lines. **(G)** Proportion of athymic nude mice that developed tumors following flank injection of *in-vivo-*R *Arid1a* KO cells overexpressing *ARID1A* or *GFP*. **(H)** Proportion of athymic nude mice that developed tumors following flank injection of *in-vivo-*R *Arid1a* Het or KO cell lines. For **(A)** through **(H)**, the proportions were compared via Fisher’s exact test. **(I)** CD4^+^ and CD8^+^ T-cell depletion in immunocompetent mice treated with anti-CD4 and anti-CD8 antibodies. Gated on FVD^-^CD45^+^CD3^+^ cells. **(J)** Tumor growth curves following flank injection of *in-vivo*-R *Arid1a* KO tumoroids into immunocompetent mice treated with control IgG, anti-CD8 antibody, or combined anti-CD4 and anti-CD8 antibodies. Data shown as mean ± SEM. Statistical comparison versus IgG control through day 28 by mixed-effects model (REML) with Geisser-Greenhouse correction measuring time × treatment interaction. **(K)** Proportion of mice that developed tumors following flank injection of *in-vivo-*R *Arid1a* KO tumoroids under the indicated T-cell depletion conditions. Proportions compared via Fisher’s exact test. **(L-N)** Flow cytometric analysis of *in-vivo-*R *Arid1a* WT and KO tumoroids 7 and 14 days after flank injection into syngeneic immunocompetent mice, measuring for infiltrated **(L)** CD45^+^ cells, **(M)** F4/80^+^CD11b^+^CD206^−^ macrophages, and **(N)** F4/80^+^CD11b^+^CD206^+^ macrophages. Data shown as mean ± SEM and compared using Welch’s t-test at each timepoint. NS, non-significant, * P < 0.05, *** P < 0.001, **** P < 0.0001.

**Supplemental Figure 4.**
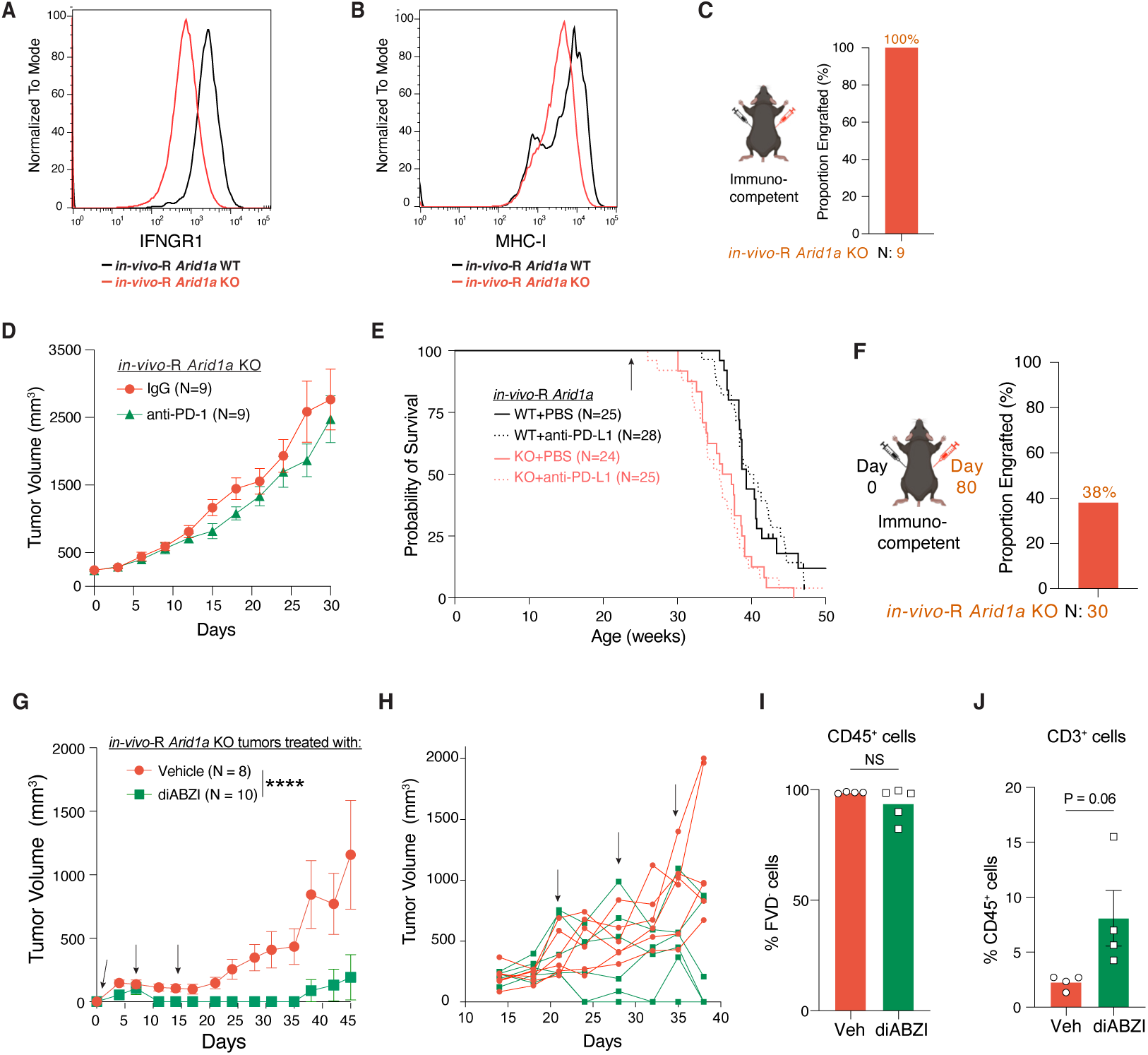
Related to Figure 4. **(A)** Representative histogram of flow cytometric analysis of cell-surface IFNGR1 expression on *in-vivo-*R *Arid1a* WT and KO tumoroids. **(B)** Representative histogram of flow cytometric analysis of cell-surface MHC-I expression on *in-vivo-*R *Arid1a* WT and KO tumoroids following IFN-γ stimulation. **(C)** Proportion of *in-vivo*-R *Arid1a* KO tumoroids that engrafted following contralateral flank co-injection with *in-vivo*-R *Arid1a* WT tumoroids in syngeneic immunocompetent mice. **(D)** Tumor growth curves of *in-vivo*-R *Arid1a* KO tumors injected into the flanks of syngeneic immunocompetent mice and then treated with control IgG or anti–PD-1 antibody. **(E)** Kaplan–Meier estimates of overall survival for *in-vivo*-R *Arid1a* WT and KO mice treated with PBS or anti–PD-L1 antibody beginning at 24 weeks of age (arrow). **(F)** Proportion of *in-vivo*-R *Arid1a* KO tumoroids that engrafted following flank injection into immunocompetent mice previously exposed to *in-vivo*-R *Arid1a* WT tumoroids. **(G)** Tumor growth curves following flank injection of *in-vivo*-R *Arid1a* KO tumoroids in mice treated with vehicle or diABZI. Treatment (arrows) was administered at days 1, 7, and 14 after tumoroid injection. Data shown as mean ± SEM. Statistical comparison of vehicle versus diABZI through day 45 by mixed-effects model (REML) with Geisser-Greenhouse correction. Asterisks denote the time × treatment interaction. **(H)** Individual tumor growth trajectories after treatment with vehicle or diABZI given at days 21, 28, and 35 after tumoroid injection (arrows indicate treatment days). **(I-J)** Flow cytometric analysis of *in-vivo*-R *Arid1a* KO tumors following intratumoral diABZI treatment, with data shown as mean ± SEM and compared with Welch’s t-test. **(I)** CD45^+^ cells as a proportion of FVD^−^ viable cells, and **(J)** CD3^+^ cells as a proportion of CD45^+^ cells. NS, non-significant, **** P < 0.0001.

**Supplemental Figure 5.**
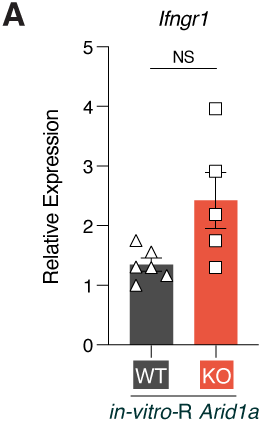
Related to Figure 5. **(A)** qPCR analysis of *Ifngr1* expression in *in-vitro-*R *Arid1a* WT and KO tumoroids. Data shown as individual data points with mean ± SEM compared with Welch’s t-test.

**Supplemental Figure 6.**
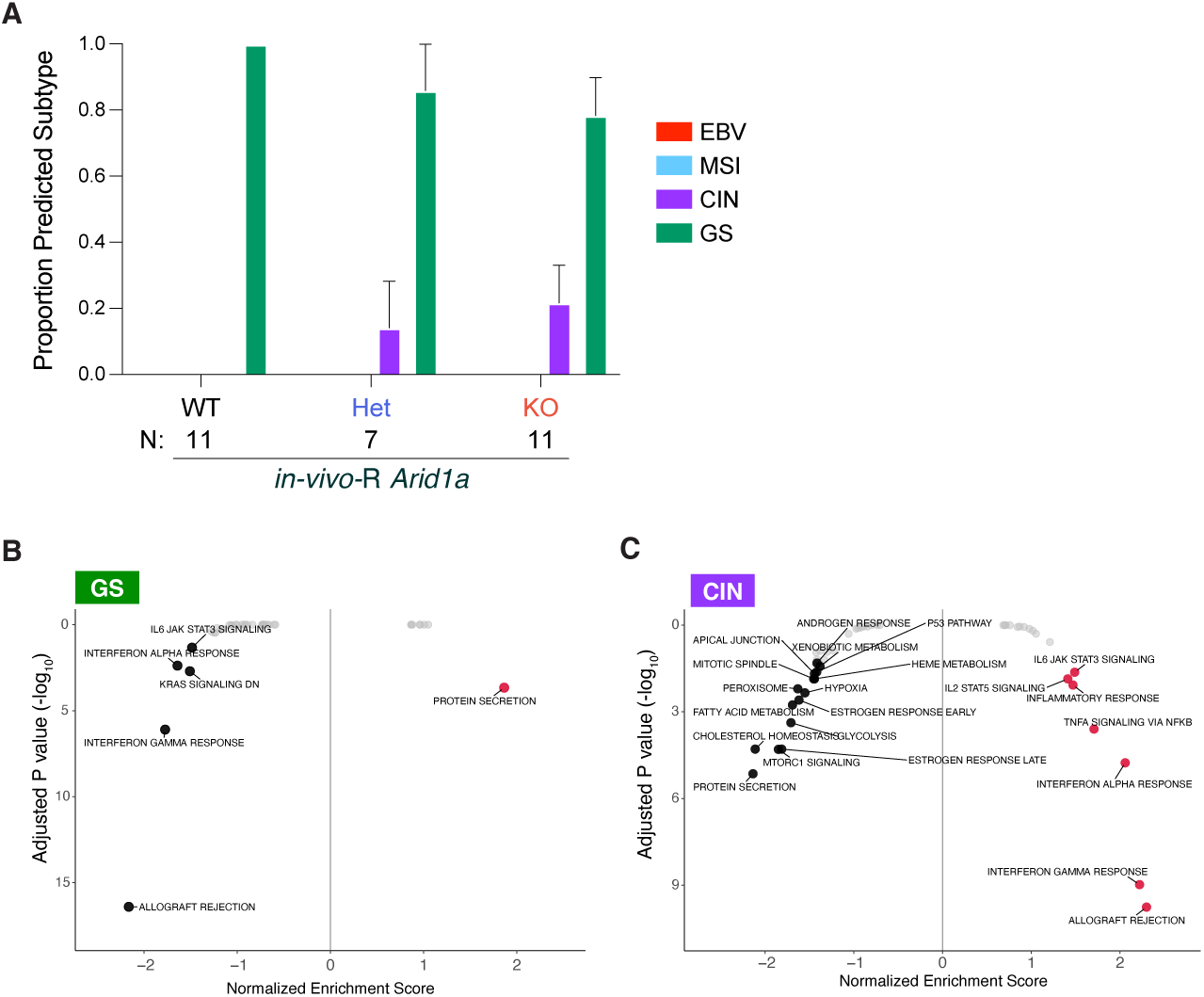
Related to Figure 6. **(A)** Mean proportion of predicted TCGA molecular subtype in gastric cancers from *in-vivo-*R *Arid1a* WT, Het, and KO mice. Data shown as mean ± SEM. (**B-C**) Gene set enrichment analysis comparing **(B)** genomically stable (GS) tumors that carried *ARID1A* LOF mutations to *ARID1A* WT samples and **(C)** chromosomal instability (CIN) tumors that carried *ARID1A* loss-of-function (LOF) mutations to *ARID1A* wild-type (WT) tumors. All significantly enriched and depleted pathways labeled.

## Notes

### Competing Interest Statement

The authors have declared no competing interest.

## REFERENCES

1. Kadoch, C., Hargreaves, D.C., Hodges, C., Elias, L., Ho, L., Ranish, J., and Crabtree, G.R. (2013). Proteomic and bioinformatic analysis of mammalian SWI/SNF complexes identifies extensive roles in human malignancy. Nat Genet 45, 592–601. 10.1038/ng.2628.

2. Sun, X.X., Chuang, J.C., Kanchwala, M., Wu, L.W., Celen, C., Li, L., Liang, H.Q., Zhang, S.Y., Maples, T., Nguyen, L.H., et al. (2016). Suppression of the SWI/SNF Component Arid1a Promotes Mammalian Regeneration. Cell Stem Cell 18, 456–466. 10.1016/j.stem.2016.03.001.

3. Shen, J., Ju, Z., Zhao, W., Wang, L., Peng, Y., Ge, Z., Nagel, Z.D., Zou, J., Wang, C., Kapoor, P., et al. (2018). ARID1A deficiency promotes mutability and potentiates therapeutic antitumor immunity unleashed by immune checkpoint blockade. Nat Med 24, 556–562. 10.1038/s41591-018-0012-z.

4. Brodeur, M.N., Dopeso, H., Zhu, Y., Longhini, A.L.F., Gazzo, A., Sun, S., Koche, R.P., Qu, R., Rosenberg, L., Hamard, P.J., et al. (2024). Interferon response and epigenetic modulation by SMARCA4 mutations drive ovarian tumor immunogenicity. Sci Adv 10, eadk4851. 10.1126/sciadv.adk4851.

5. Maxwell, M.B., Hom-Tedla, M.S., Yi, J., Li, S., Rivera, S.A., Yu, J., Burns, M.J., McRae, H.M., Stevenson, B.T., Coakley, K.E., et al. (2024). ARID1A suppresses R-loop-mediated STING-type I interferon pathway activation of anti-tumor immunity. Cell 187, 3390–3408.e3319. 10.1016/j.cell.2024.04.025.

6. Li, J., Wang, W., Zhang, Y., Cieslik, M., Guo, J., Tan, M., Green, M.D., Wang, W., Lin, H., Li, W., et al. (2020). Epigenetic driver mutations in ARID1A shape cancer immune phenotype and immunotherapy. J Clin Invest. 10.1172/JCI134402.

7. Li, N., Liu, Q., Han, Y., Pei, S., Cheng, B., Xu, J., Miao, X., Pan, Q., Wang, H., Guo, J., et al. (2022). ARID1A loss induces polymorphonuclear myeloid-derived suppressor cell chemotaxis and promotes prostate cancer progression. Nature Communications 13, 7281. 10.1038/s41467-022-34871-9.

8. Watson, P.M., DeVaux, C.A., and Freeman, K.W. (2025). Loss of ARID1A leads to a cold tumor phenotype via suppression of IFNgamma signaling. Sci Rep 15, 8716. 10.1038/s41598-025-91688-4.

9. Okamura, R., Kato, S., Lee, S., Jimenez, R.E., Sicklick, J.K., and Kurzrock, R. (2020). ARID1A alterations function as a biomarker for longer progression-free survival after anti-PD-1/PD-L1 immunotherapy. J Immunother Cancer 8. 10.1136/jitc-2019-000438.

10. Gu, Y., Zhang, P., Wang, J., Lin, C., Liu, H., Li, H., He, H., Li, R., Zhang, H., and Zhang, W. (2023). Somatic ARID1A mutation stratifies patients with gastric cancer to PD-1 blockade and adjuvant chemotherapy. Cancer Immunol Immunother 72, 1199–1208. 10.1007/s00262-022-03326-x.

11. Abou Alaiwi, S., Nassar, A.H., Xie, W., Bakouny, Z., Berchuck, J.E., Braun, D.A., Baca, S.C., Nuzzo, P.V., Flippot, R., Mouhieddine, T.H., et al. (2020). Mammalian SWI/SNF complex genomic alterations and immune checkpoint blockade in solid tumors. Cancer Immunology Research, canimm.0866.2019. 10.1158/2326-6066.Cir-19-0866.

12. Jing, H., Zhang, X., and Meng, L. (2025). Swinging the SWI/SNF complexes for cancer therapy. Trends in Pharmacological Sciences 46, 907–921. 10.1016/j.tips.2025.07.008.

13. Schreiber, R.D., Old, L.J., and Smyth, M.J. (2011). Cancer immunoediting: integrating immunity’s roles in cancer suppression and promotion. Science 331, 1565–1570. 10.1126/science.1203486.

14. Roerden, M., and Spranger, S. (2025). Cancer immune evasion, immunoediting and intratumour heterogeneity. Nat Rev Immunol 25, 353–369. 10.1038/s41577-024-01111-8.

15. The Cancer Genome Atlas Research, N. (2014). Comprehensive molecular characterization of gastric adenocarcinoma. Nature 513, 202–209. 10.1038/nature13480.

16. Gao, X., Tate, P., Hu, P., Tjian, R., Skarnes, W.C., and Wang, Z. (2008). ES cell pluripotency and germ-layer formation require the SWI/SNF chromatin remodeling component BAF250a. Proc Natl Acad Sci U S A 105, 6656–6661. 10.1073/pnas.0801802105.

17. Drost, J., van Jaarsveld, R.H., Ponsioen, B., Zimberlin, C., van Boxtel, R., Buijs, A., Sachs, N., Overmeer, R.M., Offerhaus, G.J., Begthel, H., et al. (2015). Sequential cancer mutations in cultured human intestinal stem cells. Nature 521, 43–47. 10.1038/nature14415.

18. Grzelak, C.A., Goddard, E.T., Lederer, E.E., Rajaram, K., Dai, J., Shor, R.E., Lim, A.R., Kim, J., Beronja, S., Funnell, A.P.W., and Ghajar, C.M. (2022). Elimination of fluorescent protein immunogenicity permits modeling of metastasis in immune-competent settings. Cancer Cell 40, 1–2. 10.1016/j.ccell.2021.11.004.

19. Shimada, S., Mimata, A., Sekine, M., Mogushi, K., Akiyama, Y., Fukamachi, H., Jonkers, J., Tanaka, H., Eishi, Y., and Yuasa, Y. (2012). Synergistic tumour suppressor activity of E-cadherin and p53 in a conditional mouse model for metastatic diffuse-type gastric cancer. Gut 61, 344–353. 10.1136/gutjnl-2011-300050.

20. Burclaff, J., Osaki, L.H., Liu, D., Goldenring, J.R., and Mills, J.C. (2017). Targeted Apoptosis of Parietal Cells Is Insufficient to Induce Metaplasia in Stomach. Gastroenterology 152, 762–766.e767. 10.1053/j.gastro.2016.12.001.

21. Sun, X.X., Wang, S.C., Wei, Y.L., Luo, X., Jia, Y.M., Li, L., Gopal, P., Zhu, M., Nassour, I., Chuang, J.C., et al. (2017). Arid1a Has Context-Dependent Oncogenic and Tumor Suppressor Functions in Liver Cancer. Cancer Cell 32, 574-+. 10.1016/j.ccell.2017.10.007.

22. Xu, C., Huang, K.K., Law, J.H., Chua, J.S., Sheng, T., Flores, N.M., Pizzi, M.P., Okabe, A., Tan, A.L.K., Zhu, F., et al. (2023). Comprehensive molecular phenotyping ofARID1A-deficient gastric cancer reveals pervasive epigenomic reprogramming and therapeutic opportunities. Gut. 10.1136/gutjnl-2022-328332.

23. Subramanian, A., Tamayo, P., Mootha, V.K., Mukherjee, S., Ebert, B.L., Gillette, M.A., Paulovich, A., Pomeroy, S.L., Golub, T.R., Lander, E.S., and Mesirov, J.P. (2005). Gene set enrichment analysis: A knowledge-based approach for interpreting genome-wide expression profiles. Proceedings of the National Academy of Sciences 102, 15545–15550. 10.1073/pnas.0506580102.

24. Dighe, A.S., Richards, E., Old, L.J., and Schreiber, R.D. (1994). Enhanced in vivo growth and resistance to rejection of tumor cells expressing dominant negative IFN gamma receptors. Immunity 1, 447–456. 10.1016/1074-7613(94)90087-6.

25. Shankaran, V., Ikeda, H., Bruce, A.T., White, J.M., Swanson, P.E., Old, L.J., and Schreiber, R.D. (2001). IFNgamma and lymphocytes prevent primary tumour development and shape tumour immunogenicity. Nature 410, 1107–1111. 10.1038/35074122.

26. Chen, D.S., and Mellman, I. (2017). Elements of cancer immunity and the cancer-immune set point. Nature 541, 321–330. 10.1038/nature21349.

27. Bianchi, A., De Castro Silva, I., Deshpande, N.U., Singh, S., Mehra, S., Garrido, V.T., Guo, X., Nivelo, L.A., Kolonias, D.S., Saigh, S.J., et al. (2023). Cell-Autonomous Cxcl1 Sustains Tolerogenic Circuitries and Stromal Inflammation via Neutrophil-Derived TNF in Pancreatic Cancer. Cancer Discov 13, 1428–1453. 10.1158/2159-8290.Cd-22-1046.

28. Adrover, J.M., Han, X., Sun, L., Fujii, T., Sivetz, N., Daßler-Plenker, J., Evans, C., Peters, J., He, X.-Y., Cannon, C.D., et al. (2025). Neutrophils drive vascular occlusion, tumour necrosis and metastasis. Nature 645, 484–495. 10.1038/s41586-025-09278-3.

29. Mellman, I., Chen, D.S., Powles, T., and Turley, S.J. (2023). The cancer-immunity cycle: Indication, genotype, and immunotype. Immunity 56, 2188–2205. 10.1016/j.immuni.2023.09.011.

30. Dougan, M., Dranoff, G., and Dougan, S.K. (2019). GM-CSF, IL-3, and IL-5 Family of Cytokines: Regulators of Inflammation. Immunity 50, 796–811. 10.1016/j.immuni.2019.03.022.

31. Gocher, A.M., Workman, C.J., and Vignali, D.A.A. (2022). Interferon-γ: teammate or opponent in the tumour microenvironment? Nature Reviews Immunology 22, 158–172. 10.1038/s41577-021-00566-3.

32. Lanng, K.R.B., Lauridsen, E.L., and Jakobsen, M.R. (2024). The balance of STING signaling orchestrates immunity in cancer. Nat Immunol 25, 1144–1157. 10.1038/s41590-024-01872-3.

33. Ramanjulu, J.M., Pesiridis, G.S., Yang, J., Concha, N., Singhaus, R., Zhang, S.-Y., Tran, J.-L., Moore, P., Lehmann, S., Eberl, H.C., et al. (2018). Design of amidobenzimidazole STING receptor agonists with systemic activity. Nature 564, 439–443. 10.1038/s41586-018-0705-y.

34. Ayers, M., Lunceford, J., Nebozhyn, M., Murphy, E., Loboda, A., Kaufman, D.R., Albright, A., Cheng, J.D., Kang, S.P., Shankaran, V., et al. (2017). IFN-gamma-related mRNA profile predicts clinical response to PD-1 blockade. J Clin Invest 127, 2930–2940. 10.1172/JCI91190.

35. Wang, S.C., Nassour, I., Xiao, S., Zhang, S., Luo, X., Lee, J., Li, L., Sun, X., Nguyen, L.H., Chuang, J.C., et al. (2019). SWI/SNF component ARID1A restrains pancreatic neoplasia formation. Gut 68, 1259–1270. 10.1136/gutjnl-2017-315490.

36. Okada, M., Yamasaki, S., Nakazato, H., Hirahara, Y., Ishibashi, T., Kawamura, M., Shimizu, K., and Fujii, S.I. (2024). ARID1A-Deficient Tumors Acquire Immunogenic Neoantigens during the Development of Resistance to Targeted Therapy. Cancer Res 84, 2792–2805. 10.1158/0008-5472.CAN-23-2846.

37. Shi, W., Wang, Y., Xu, C., Li, Y., Ge, S., Bai, B., Zhang, K., Wang, Y., Zheng, N., Wang, J., et al. (2023). Multilevel proteomic analyses reveal molecular diversity between diffuse-type and intestinal-type gastric cancer. Nat Commun 14, 835. 10.1038/s41467-023-35797-6.

38. Lauren, P. (1965). The Two Histological Main Types of Gastric Carcinoma: Diffuse and So-Called Intestinal-Type Carcinoma. An Attempt at a Histo-Clinical Classification. Acta Pathol Microbiol Scand 64, 31–49. 10.1111/apm.1965.64.1.31.

39. Lee, J.H., Chang, K.K., Yoon, C., Tang, L.H., Strong, V.E., and Yoon, S.S. (2018). Lauren Histologic Type Is the Most Important Factor Associated With Pattern of Recurrence Following Resection of Gastric Adenocarcinoma. Ann Surg 267, 105–113. 10.1097/sla.0000000000002040.

40. Jiménez Fonseca, P., Carmona-Bayonas, A., Hernández, R., Custodio, A., Cano, J.M., Lacalle, A., Echavarria, I., Macias, I., Mangas, M., Visa, L., et al. (2017). Lauren subtypes of advanced gastric cancer influence survival and response to chemotherapy: real-world data from the AGAMENON National Cancer Registry. British Journal of Cancer 117, 775–782. 10.1038/bjc.2017.245.

41. Hoyos, D., Zappasodi, R., Schulze, I., Sethna, Z., de Andrade, K.C., Bajorin, D.F., Bandlamudi, C., Callahan, M.K., Funt, S.A., Hadrup, S.R., et al. (2022). Fundamental immune-oncogenicity trade-offs define driver mutation fitness. Nature 606, 172–179. 10.1038/s41586-022-04696-z.

## SUPPLEMENTAL REFERENCES

1 FastQC: A Quality Control tool for High Throughput Sequence Data.

2 Trim Galore!

3 Li, H. Aligning sequence reads, clone sequences and assembly contigs with BWA-MEM. arXiv 1303.3997 (2013).

4 Li, H. et al. The Sequence Alignment/Map format and SAMtools. Bioinformatics 25, 2078–2079 (2009).

5 Picard toolkit (2019).

6 Barnett, D. W., Garrison, E. K., Quinlan, A. R., Strömberg, M. P. & Marth, G. T. BamTools: a C++ API and toolkit for analyzing and managing BAM files. Bioinformatics 27, 1691–1692 (2011).

7 Amemiya, H. M., Kundaje, A. & Boyle, A. P. The ENCODE Blacklist: Identification of Problematic Regions of the Genome. Scientific Reports 9, 9354 (2019).

8 Ramírez, F. et al. deepTools2: a next generation web server for deep-sequencing data analysis. Nucleic Acids Research 44, W160–W165 (2016).

9 Zhang, Y. et al. Model-based Analysis of ChIP-Seq (MACS). Genome Biology 9, R137 (2008).

10 Orchard, P., Kyono, Y., Hensley, J., Kitzman, J. O. & Parker, S. C. J. Quantification, Dynamic Visualization, and Validation of Bias in ATAC-Seq Data with ataqv. Cell Systems 10, 298–306.e294 (2020).

11 DiffBind: Differential binding analysis of ChIP-Seq peak data (2011).

12 Ross-Innes, C. S. et al. Differential oestrogen receptor binding is associated with clinical outcome in breast cancer. Nature 481, 389–393 (2012).

13 Zhu, L. J. in Tiling Arrays: Methods and Protocols (eds T. L. Lee & A. C. Shui Luk) 105–124 (Humana Press, 2013).

14 Zhu, L. J. et al. ChIPpeakAnno: a Bioconductor package to annotate ChIP-seq and ChIP-chip data. BMC Bioinformatics 11, 237 (2010).

15 Hahne, F. & Ivanek, R. in Statistical Genomics: Methods and Protocols (eds E. Mathé & S. Davis) 335–351 (Springer, 2016).

16 Dobin, A. et al. STAR: ultrafast universal RNA-seq aligner. Bioinformatics 29, 15–21 (2013).

17 Liao, Y., Smyth, G. K. & Shi, W. featureCounts: an efficient general purpose program for assigning sequence reads to genomic features. Bioinformatics 30, 923–930 (2014).

18 Love, M. I., Huber, W. & Anders, S. Moderated estimation of fold change and dispersion for RNA-seq data with DESeq2. Genome Biol 15, 550 (2014). 10.1186/s13059-014-0550-8

19 Korotkevich, G., Sukhov, V., Budin, N., Shpak, B., Artyomov, M.N., & Sergushichev, A.

20 Liberzon, A., Birger, C., Thorvaldsdóttir, H., Ghandi, M., Mesirov, J.P., & Tamayo, P.

21 Patro, R., Duggal, G., Love, M. I., Irizarry, R. A. & Kingsford, C. Salmon provides fast and bias-aware quantification of transcript expression. Nat Methods 14, 417–419 (2017). 10.1038/nmeth.4197

22 Soneson, C., Love, M. I. & Robinson, M. D. Differential analyses for RNA-seq: transcript-level estimates improve gene-level inferences. F1000Res 4, 1521 (2015). 10.12688/f1000research.7563.2

23 Durinck, S., Spellman, P. T., Birney, E. & Huber, W. Mapping identifiers for the integration of genomic datasets with the R/Bioconductor package biomaRt. Nat Protoc 4, 1184–1191 (2009). 10.1038/nprot.2009.97

24 Cerami, E. et al. The cBio cancer genomics portal: an open platform for exploring multidimensional cancer genomics data. Cancer Discov 2, 401–404 (2012). 10.1158/2159-8290.CD-12-0095

25 Gao, J. et al. Integrative analysis of complex cancer genomics and clinical profiles using the cBioPortal. Sci Signal 6, pl1 (2013). 10.1126/scisignal.2004088

26 Tibshirani, R., Hastie, T., Narasimhan, B. & Chu, G. Diagnosis of multiple cancer types by shrunken centroids of gene expression. Proc Natl Acad Sci U S A 99, 6567–6572 (2002). 10.1073/pnas.082099299

27 Ritchie, M. E. et al. limma powers differential expression analyses for RNA-sequencing and microarray studies. Nucleic Acids Res 43, e47 (2015). 10.1093/nar/gkv007

28 Rosenberg, A. B. et al. Single-cell profiling of the developing mouse brain and spinal cord with split-pool barcoding. Science 360, 176–182 (2018). 10.1126/science.aam8999

29 Carlson, M. Vol. 3.16.0 (Bioconductor, 2022).

30 Hao, Y. et al. Dictionary learning for integrative, multimodal and scalable single-cell analysis. Nat Biotechnol 42, 293–304 (2024). 10.1038/s41587-023-01767-y

31 Hafemeister, C. & Satija, R. Normalization and variance stabilization of single-cell RNA-seq data using regularized negative binomial regression. Genome Biol 20, 296 (2019). 10.1186/s13059-019-1874-1

32 Choudhary, S. & Satija, R. Comparison and evaluation of statistical error models for scRNA-seq. Genome Biol 23, 27 (2022). 10.1186/s13059-021-02584-9

33 Ahlmann-Eltze, C. & Huber, W. glmGamPoi: fitting Gamma-Poisson generalized linear models on single cell count data. Bioinformatics 36, 5701–5702 (2021). 10.1093/bioinformatics/btaa1009

34 Korsunsky, I. et al. Fast, sensitive and accurate integration of single-cell data with Harmony. Nat Methods 16, 1289–1296 (2019). 10.1038/s41592-019-0619-0

35 McInnes, L., Healy, J. & Melville, J. UMAP: Uniform Manifold Approximation and Projection for Dimension Reduction. arXiv (2018). 10.48550/arXiv.1802.03426

36 Blondel, V. D., Guillaume, J.-L., Lambiotte, R. & Lefebvre, E. Fast unfolding of communities in large networks. Journal of Statistical Mechanics: Theory and Experiment 2008, P10008 (2008). 10.1088/1742-5468/2008/10/p10008

